# SweepLink: Joint Inference of Demography and Linked Selection from Time-series Data

**DOI:** 10.64898/2026.09.02.748944

**Authors:** Ekaterina Noskova, Madleina Caduff, Andreas Fueglistaler, Anna Parker, Christoph Leuenberger, Daniel Wegmann

**Affiliations:** Department of Biology, University of Fribourg, 1700 Fribourg, Switzerland; Swiss Institute of Bioinformatics, 1700 Fribourg, Switzerland; Institute of Ecology and Evolution, University of Edinburgh, EH9 3DW Edinburgh, United Kingdom; Swiss Tropical Institute, 4123 Allschwil, Switzerland; Department of Mathematics, University of Fribourg, 1700 Fribourg, Switzerland

## Abstract

Genome-wide time-series data, i.e. allele frequency trajectories tracked across multiple sampling times, are among the richest sources of information for inferring selection. Beyond a beneficial allele’s own rise in frequency, such data capture how it drags nearby loci upward via linkage, an effect known as genetic hitch-hiking. Yet most existing tools are single-locus, treating loci independently: they infer site-specific selection coefficients in isolation, then rely on ad hoc window statistics to account for hitch-hiking. Many existing tools further require a predefined population size, or scale poorly when jointly inferring selection and demography, and their power is highly sensitive to a significance threshold. To address these shortcomings, we here present `SweepLink`, a two-layer Hidden Markov Model that overcomes these limitations by jointly inferring demography and linked selection genome-wide: a spatial layer captures correlations between neighboring selection coefficients, coupled with a temporal Wright-Fisher diffusion layer. As we show with extensive simulations, this setup pushes drift-driven false signals toward neutrality while reinforcing loci that receive support from neighbouring loci, thereby increasing the sensitivity for weak and moderate selection, while matching the power of existing tools to detect strong selection. These simulations further show that `SweepLink` yields confident posteriors that remain stable at maximal significance, removing the need for arbitrary thresholds. We applied `SweepLink` to ancient DNA time-series data from the British population, previously analysed with a single-locus tool. `SweepLink` recovers four of the previously reported signals (LCT, SLC45A2, DHCR7, HERC2), and partially recovers the MHC/HLA signal. It also identifies additional candidate regions, including DPYD, FADS1/2 and OAS1, missed by the prior scan but supported by independent studies.

## 1 Introduction

Selective signals provide critical information about the adaptive processes that have shaped biological populations. The emergence of ancient DNA (aDNA) has significantly boosted the power of these analyses by providing time-series data. These temporal data allow for the direct observation of how allele frequencies change over generations. Beneficial alleles under positive selection are expected to increase in frequency and eventually reach fixation, while deleterious alleles under negative selection are expected to decrease the frequency and eventually be lost. Since the speed of these processes is modulated by the strength of selection, allele frequency trajectories are thus directly informative regarding both the strength and mode of selection.

Allele frequency trajectories are, however, also affected by genetic drift. Consequently, the inference of selection is intrinsically linked to demography, and failing to account for demographic history is well known to bias selection estimates [Jensen et al., 2005, Nielsen et al., 2005, Mughal and DeGiorgio, 2019]. To mitigate this bias in case of time-series data, methods generally model genetic drift using the Wright-Fisher process using a temporal HMM layer [e.g. Malaspinas et al., 2012, Ferrer-Admetlla et al., 2016, Mathieson and Terhorst, 2022, Cheng and Steinrücken, 2025]. Recent tools, such as `bmws` [Mathieson and Terhorst, 2022] and `diplo-locus` [Cheng and Steinrücken, 2025], successfully scale this framework to genome-wide data by utilizing highly efficient maximum likelihood inference and requiring a predefined effective population size. Consequently, they provide only point estimates and remain vulnerable to demographic misspecification. To overcome the dependence on predefined demographic parameters, several methods employ Bayesian inference to jointly estimate both population size and selection coefficients from multi-locus data sets. As this is computationally demanding, early approaches were simulation-based and used Approxmate Bayesian Computation (ABC) for inference [Foll et al., 2015], an approach that can scale efficiently to large data sets [Kousathanas et al., 2016]. As an alternative, `ApproxWF` [Ferrer-Admetlla et al., 2016] introduced an efficient approximation to model the Wright-Fisher process on a discrete allele frequency grid, which can be fully evaluated using a temporal HMM approach. `ApproxWF` was found to be particularly poweful at identifying selected loci in a recent comparison [Anchieri et al., 2026], and has also been successfully applied to ancient DNA time series data [e.g. Burger et al., 2020].

Despite their different strategies for handling demography and computational scalability, all of these methods assume that loci evolve independently and thereby ignore the broader genomic context. This assumption fails to account for the “hitch-hiking effect”, where selection acting on a causal variant also influences the frequencies of nearby loci [Barton, 2000]. It has been demonstrated through extensive simulations that ignoring such linked selection leads to significantly biased estimates of selection coefficients, which can be overcome by modelling linked sites explicitly [He et al., 2020]. However, the explicit model introduced in [He et al., 2020] accounts for a single pair of loci, and modelling the linkage structure at a large number of sites has proven computationally demanding. Terhorst et al. [2015], for instance, introduced a method to jointly analyze many linked sites by approximating the multi-locus Wright-Fisher model using a Gaussian process that captures correlations between sites through its covariance structure. However, the runtime scales quadratically with the number of loci, confining any inference to small local windows rather than the full genomic landscape. A related strategy estimates this covariance structure directly from allele-frequency trajectories without requiring phased haplotypes [Li and Barton, 2023], though at a scale (tens of loci) still order of magnitude smaller than genome-wide analysis. Rather than modeling the linkage structure explicitly, `timesweeper` [Whitehouse and Schrider, 2023] trains a machine learning classifier on data simulated data. The computational burden remains high, however, as the training requires extensive forward-in-time simulations, which are impractical for species with large effective population sizes (such as humans) and large genomic windows. Another alternative is `CLUES2` [Vaughn and Nielsen, 2024], which leverages Ancestral Recombination Graphs (ARGs) to account for linkage, but the computational cost and data requirements to infer these remain a major bottleneck.

To address these limitations, we present `SweepLink`, a novel method for joint inference of demography and linked selection from time-series allele count data that captures correlations between loci through an additional genome-wide layer within its HMM framework on locus-specific selection coefficients. As our experiments on simulated data demonstrate, `SweepLink` had higher sensitivity to weak selection compared to standard single-locus tools, with a notable reduction of false-positives, despite remaining computationally tractable and scaling to full genome analyses. Moreover, by pooling evidence of selection across the genomic landscape, our method produces more confident posterior probabilities, eliminating the need for arbitrary empirical thresholds to distinguish true from false selective signals. Applying `SweepLink` to time-series data from British populations, we recover more known targets of historical adaptation than single-locus methods and identify several novel candidate regions.

## 2 Materials and Methods

### 2.1 Methodological Overview

`SweepLink` is a probabilistic modelling framework designed to jointly infer demographic parameters and linked selection from time-series allele frequency data. The method is structured as a two-layer Hidden Markov Model (HMM), illustrated in Figure 1A. The first **spatial layer** models selection coefficients along the genome and accounts for the correlation between adjacent loci through a ladder-type transition matrix *Q*. Its parameters constrain adjacent loci to share similar selection coefficients, while assuming that the vast majority of the genome evolves neutrally, thereby suppressing isolated false-positive signals caused by genetic drift and pooling signals from neighbouring loci into distinct high-confidence selection peaks (Figure 1B). The second **temporal layer** incorporates a diffusion approximation of the Wright-Fisher model to calculate the likelihood of the evolutionary parameters. It thus resembles single-locus methods, such as `ApproxWF` [Ferrer-Admetlla et al., 2016] or `diplo-locus` [Cheng and Steinrücken, 2025], which also utilize the Wright-Fisher model within an HMM framework.

**Figure 1.**
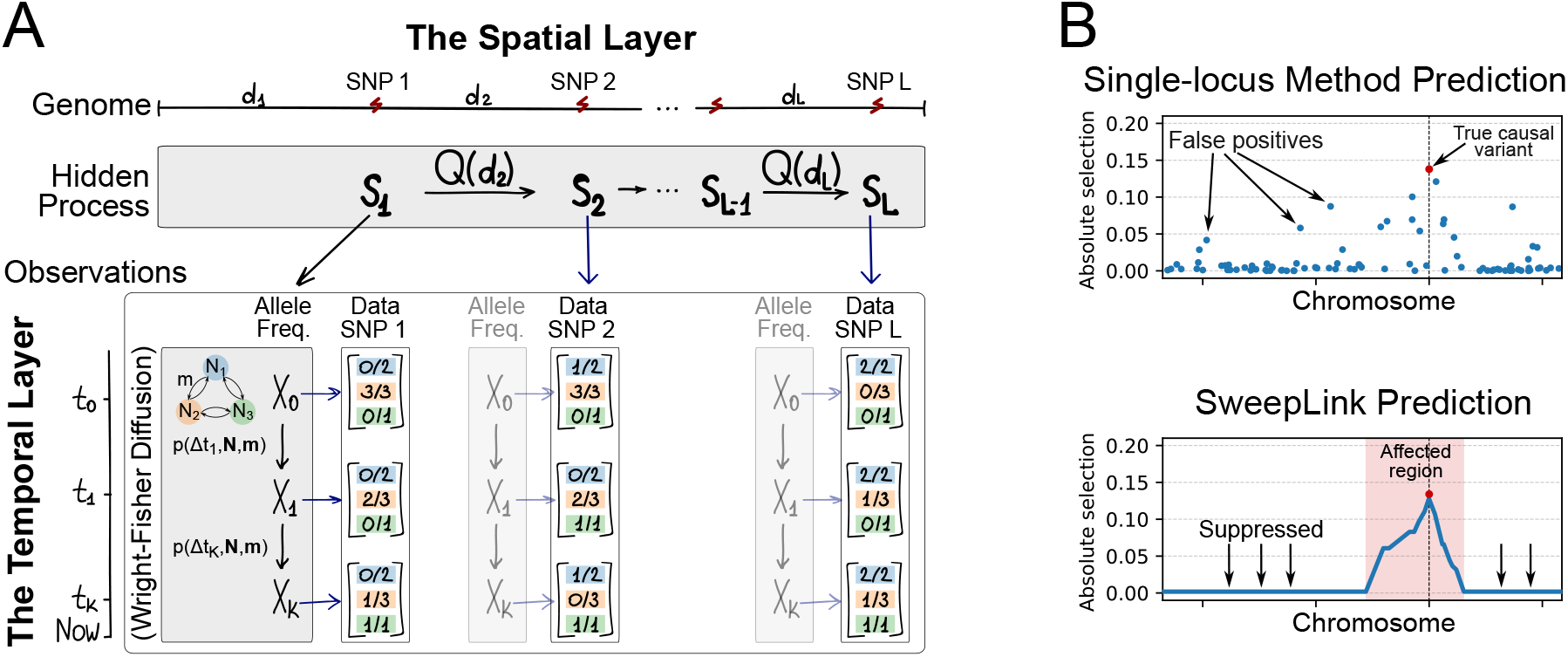
Overview of the SweepLink Two-Layer HMM architecture. **(A)** The model framework. The spatial layer (top, horizontal) models selection coefficients as a hidden Markov process (**S**_*l*_) along the genome, with transitions evaluated by distance *d*_*l*_. The temporal layer (bottom, vertical) utilizes a Wright-Fisher diffusion process to calculate the likelihood of observed allele counts. These temporal probabilities are jointly parametrized by locus-specific selection (**S**_*l*_), genome-wide population sizes (**N**), and migration rates (**m**). **(B)** Demonstration of selection inference for one population. Unlike single-locus methods (top) that are susceptible to noise from genetic drift, `SweepLink` (bottom) uses spatial linkage to suppress false-positives and pool true signals into a confident regional peak (red shading).

This joint HMM framework allows us to use the Metropolis-Hastings Markov Chain Monte Carlo (MCMC) method to infer the parameters of the transition matrix *Q*, the selection coefficients **S**_*l*_ along the genome, and the demographic parameters (**N, m**) directly from the genetic data.

### 2.2 Underlying Demographic Model and Time-Series Data

We first formalize the notation of the input genetic data and the baseline demographic assumptions. Although this study mainly focuses on the case of a single population, we here develop and implement the framework for an island model, with each population *p* maintaining a constant effective size of 2*N*_*p*_ haploid individuals and continuous migration between all pairs of populations via a constant migration matrix *m*, of which the entry *m*_*pq*_ denotes the continuous migration rate from population *q* into population *p*. However, we note that the framework can be readily extended to account for changes in population sizes and migration rates, as we will discuss below.

Consider a dataset containing *L* biallelic variants (SNPs) across the genome. The physical distance between locus *l* and locus *l* − 1 is denoted as *d*_*l*_. We have samples at *K* + 1 distinct historical time points defined as *{t*_0_ *< t*_1_ *<* … *< t*_*K*_*}*. Genetic data is presented for *P* populations, with *n*_*p,k*_ diploid individuals are sampled for population *p* at time point *t*_*k*_. For each locus *l* ∈ *{*1, …, *L}* at a given time point *t*_*k*_, the genetic data consists of observed allele counts. We denote this data as 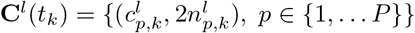, where 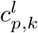 refers to the number of observed derived alleles and 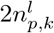 is the total number of haploid samples covering that specific locus. We note that due to potential missing data at individual sites, 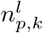 can be less or equal to the number of diploid samples *n*_*p,k*_.

### 2.3 The Spatial Layer: Genome-wide HMM for Linked Selection

When a beneficial allele emerges in a population and increases in frequency, the allele trajectories of nearby segregating loci are also affected [Barton, 2000]. If the causal beneficial allele is in linkage disequilibrium (LD) with the derived allele of a neighbouring locus, that passenger allele will hitch-hike and correspondingly increase in frequency too. Conversely, if the beneficial variant is linked to the ancestral allele, the derived allele will decrease in frequency, generating an allele trajectory that mimics the signature of negative selection. Under the Wright-Fisher model, if a focal allele experiences a selection coefficient *s*, the corresponding alternative allele experiences an inverse selective pressure defined as:

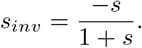

Recombination breaks down this linkage disequilibrium. As the physical distance from the causal variant increases, this breakdown of linkage causes the population-level data to show a progressively lower selection coefficient at neighbouring loci. As a result, the selection footprint forms a distinct peak, characterized by high absolute selection coefficients near the causal variant that decline toward neutrality farther away on both sides. Importantly, this pattern depends on the absolute magnitude of selection, as the directional sign (*s* or *s*_*inv*_) simply reflects the initial linkage phase of the associated alleles. The primary objective of the spatial layer is to capture this hitch-hiking effect and genetic linkage between loci.

We model the locus-specific selection coefficients as a hidden Markov process **S**(*l*), where the state **S**(*l*) is a vector (*S*_1_(*l*), …, *S*_*P*_ (*l*)) representing the population-specific selection coefficients. For simplicity, We assume each *S*_*p*_(*l*) evolves as an independent Markov process. Therefore, we omit the subscript *p* for the remainder of this section and describe the process generically for *S*(*l*), but note that more complicated models could be envisioned.

To represent selection magnitudes, we define a discrete grid of absolute values *G* = {0, *s*_1_, *s*_2_, …, *s*_*M*_*}*. Using the inverse selection definition described above, we extend this positive grid to include corresponding negative values:

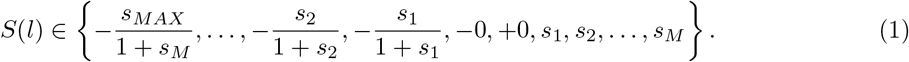

Two neutral states (− 0, +0) are explicitly retained to maintain balance in the MCMC proposal mechanism (see below).

The state *S*(*l*) is decomposed into a direction component *D*(*l*) ∈ *{*+1, −1*}* and an absolute magnitude component *Z*(*l*) ∈ *G*, as:

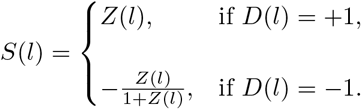

Components *D*(*l*) and *Z*(*l*) are modelled as two independent Markov chains. Since the sign on the linkage *D*(*l*) purely depends on the polarization of the alleles, its transition probabilities are uniformly equal across the genome and we have 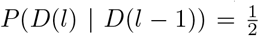. The absolute value of selection *Z*(*l*) lies within grid *G* = {0, *s*_1_, *s*_2_, …, *s*_*M*_*}*. The transition matrix of this Markov model is inspired by Galimberti et al. [2020] and defined as:

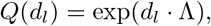

with elements [*Q*(*d*_*l*_)]_*ij*_ denoting the probability of going from state *s*_*i*_ at locus *l* − 1 to state *s*_*j*_ at locus *l* with distance *d*_*l*_ between those loci, either in physical or in recombination space. The Λ is (*M* + 1) *×* (*M* + 1) ladder-type generation matrix defined as:

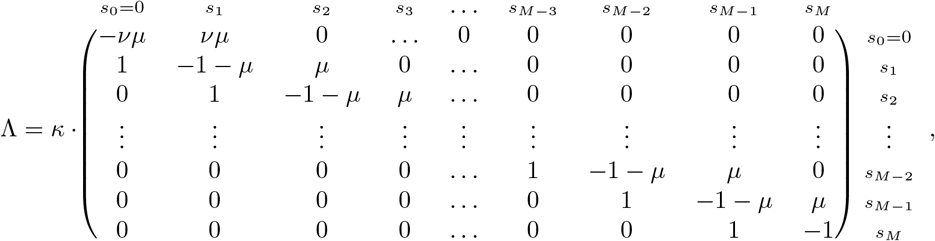

This tridiagonal structure restricts instantaneous transitions to adjacent states on the selection grid, preventing biologically implausible jumps between extreme selection values over short distances. The transition matrix *Q* is hence defined by three parameters: *κ, ν* and *µ*. The positive scaling *κ* measures the strength of correlation between loci. The first row of matrix Λ (the attractor) corresponds to neutrality and has a specific parameter *ν* ∈ [0, 1] that defines the probability to leave the neutral state. The third parameter *µ* ∈ [0, 1] models the rate of moving towards higher values of selection up to *s*_*M*_ .

Assuming the absolute magnitude *Z*(*l*) and linkage direction *D*(*l*) evolve as independent Markov processes, the transition probability for the fully signed state *S*(*l*) factors into the product of its components:

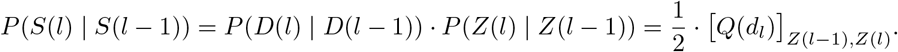

Therefore, the probability of a proposed sequence of selection coefficients **S** = (**S**(1), …, **S**(*L*)) given matrix *Q* is proportional to:

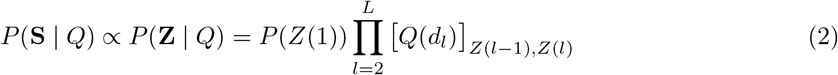

where the probability of the initial state *P* (*Z*(1)) operates as the stationary distribution of the generator matrix Λ.

### 2.4 The Temporal Layer: Hidden Diffusion Model

The temporal layer produces the emission probabilities *P* (**C**^*l*^ | **S**(*l*) = **s**) for the spatial layer at each locus *l*. The layer is defined by the demographic parameters, here the population sizes **N** and migration rates **m**. Unlike the locus-specific selection coefficients defined by the spatial layer, the demographic parameters are shared globally across all loci. For simplicity, we will neglect the notation of locus *l* in this section.

To compute the likelihood of observed allele counts **C**(*t*) at a locus given demographic parameters (***N***, ***m, s***), we apply an HMM framework in which the hidden states **X**(*t*) are true population allele frequencies evolving as a Wright–Fisher diffusion, and the emissions are binomial samples at each time point:

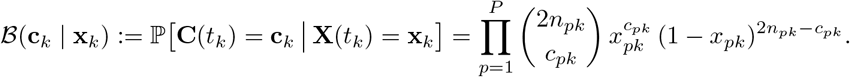

Briefly, Wright-Fisher diffusion is a continuous-time continuous state Markov process. Its transition probability density *p*(*τ*, **x, y**) – the probability of moving from frequency **x** at time *t* to frequency **y** at time *t* + *τ* – satisfies the Kolmogorov forward (Fokker-Planck) equation:

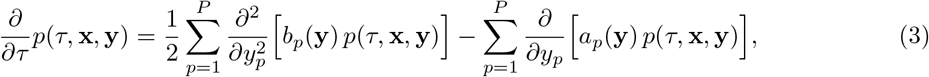

where infinitesimal mean (drift) *a*_*p*_(**y**) and variance (diffusion) *b*_*p*_(**y**) are defined by demographic parameters ***N***, ***m*** and ***s***. See Supplementary Notes for more details.

Crucially, formulating the temporal layer as a continuous diffusion process allows us to evaluate the likelihood of the time-series data using an efficient forward pass algorithm as in He et al. [2020]. We define the forward variable *α*_*k*_(*τ*, **y**) as the joint probability of the hidden frequency state at time *t*_*k*_ + *τ* and all observed data up to time *t*_*k*_:

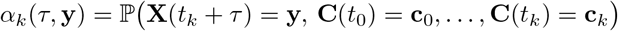

It can be shown that the recursive update of the forward variable:

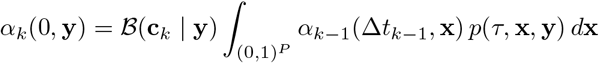

can be replaced by the the forward Kolmogorov equation 3:

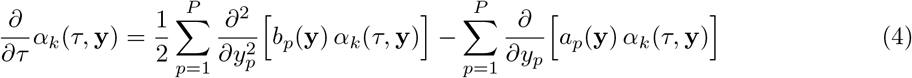

subject to the initial condition:

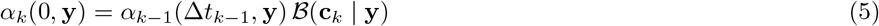

Consequently, we can iteratively evaluate the forward pass from *k* = 0 to *K* by numerically solving this partial differential equation 4 across sequential time intervals. To establish the base case for this induction at the first sampling time point (*k* = 0), the recursion is initialized using the prior density *α*(**y**), such that *α*_0_(0, **y**) = *α*(**y**)*B*(**c**_0_ | **y**). For all subsequent time points, the final solution from the preceding interval, *α*_*k*−1_(Δ*t*_*k*−1_, **y**), is substituted into Equation 5 to continuously initialize the next numerical integration.

Once the forward pass reaches the final sampling time point *t*_*K*_, the total likelihood of the model is obtained by integrating the final forward variable *α*_*K*_:

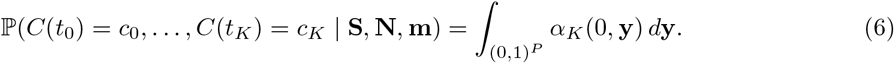

Because evaluating these multidimensional partial differential equations analytically is broadly intractable, we discretize the state space and solve the forward operator numerically using the Chang-Cooper numerical scheme Chang and Cooper [1970], Pareschi and Zanella [2018] and alternating direction method Press [2007] (for more details see Supplementary Note X). We implemented several previously proposed types of grids for the relative allele frequency space [Gutenkunst et al., 2009, Cheng and Steinrücken, 2025, Malaspinas et al., 2012, Ferrer-Admetlla et al., 2016] with the quadratic grid providing the most robust performance in empirical tests (see Supplementary Note X and Figure SX).

### 2.5 Bayesian Inference and Parameter Estimation

To infer the evolutionary history from the time-series data, we employ a Bayesian framework to estimate the joint posterior distribution of the model parameters. We first summarize the complete set of parameters estimated within the `SweepLink` model. The spatial layer introduces **S** = {**S**(1), **S**(2), …, **S**(*L*)} as the hidden sequence of locus-specific selection states along the genome, alongside the parameters ***κ, ν, µ*** that define the transition matrix *Q*. The parameter space is subsequently completed by the global demographic variables **N** and **m** derived from the temporal layer. Given the observed allele count data **C**, the posterior distribution for the parameter set ***θ*** = (**S, *κ, ν, µ*, N, m**) is proportional to:

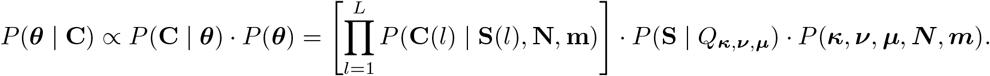

Here, the locus-specific emission probabilities *P* (**C**(*l*) | **S**(*l*), **N, m**) are defined by Equation 6, the probability of the full vector of selection coefficients *P* (**S** | *Q*_***κ***,***ν***,***µ***_) is defined by Equation 2, and *P* (***κ, ν, µ, N***, ***m***) evaluates as the combined product of the independent parameter priors.

We utilize a Markov chain Monte Carlo (MCMC) algorithm to draw representative samples directly from this posterior distribution. Specifically, we execute a Metropolis-Hastings sampling scheme [Metropolis et al., 1953, Hastings, 1970] that iteratively proposes new parameter values and hidden states, accepting or rejecting them based on the evaluated ratios of this unnormalized posterior distribution.

To translate the resulting MCMC trace into quantitative evidence for selection at any given locus, we approximate the marginal posterior probabilities directly from the parameter samples. Let *W* be the total number of MCMC samples. For any specific selection coefficient *s* on the fully signed discrete grid, its estimated posterior probability is calculated as its sample frequency, utilizing a smoothing correction defined as *X*_*s*_*/*(*W* + 1), where *X*_*s*_ is the number of MCMC samples yielding state *s*. We then calculate the posterior probability *P* (*s >* 0) of positive selection, and probability *P* (*s <* 0) of negative selection by summing these estimates across all strictly positive and strictly negative states on the grid. Finally, to summarize the model’s overall confidence in directional selection, we define our primary classification score as:

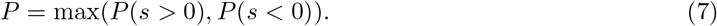

This value serves as `SweepLink` ‘s primary statistic to distinguish between neutral variants and selective sweeps.

### 2.6 Implementation

The proposed model and the Bayesian inference scheme are implemented in an easy-to-use C++ program `SweepLink`, available at https://bitbucket.org/wegmannlab/sweeplink/. The input file is analogous to that of `ApproxWF` [Ferrer-Admetlla et al., 2016] and contains allele counts per time point and population. These allele counts can be estimated from genotype likelihoods using `ATLAS` [Link et al., 2017]. Alternatively, a provided Python scripts extracts these counts from VCF files. This script also post-processes `SweepLink` output, including plotting of the results. Detailed documentation and a tutorial can be found at https://sweeplink.readthedocs.io. We note that while the code provided works with an island model of arbitrary size, computation time grows quadratically with the number of populations and we thus focus here primarily on the case of a single population.

### 2.7 Benchmarking Against Competitor Single-Locus Tools

#### Simulated Benchmark Dataset

To perform a benchmark analysis of `SweepLink`, we generated ten short chromosomes of length 0.1 Mb each using the forward-in-time simulator SLiM v4 [Haller and Messer, 2023]. Half of the chromosomes contained only neutral mutations, while the remaining five chromosomes had a hard selective sweep introduced at the middle position, with one chromosome for each of the selection coefficients 0.01, 0.02, 0.03, 0.04, or 0.05. Population parameters were chosen to resemble realistic human populations: population size was set to *N* = 10,000 individuals [Takahata, 1993], the mutation rate to *µ*_*aA*_ = 10^−8^ per site per generation [Kong et al., 2012], and the recombination rate to *r* = 10^−8^ per site per generation [Kong et al., 2002].

Sampling was performed once the selected allele reaches a frequency of 30% for 160 generations at *K* = 11 consecutive time points spaced by Δ*t* = 16 generations apart. At each time point, genetic data for *n* = 25 diploid individuals were sampled. This sampling regime roughly matches the real human data analysed in this study, which includes samples dating to the last 4,000 years. We generated a total of 100 independent replicates of this setup, yielding a total dataset of 500 independent selective sweeps (100 for each selection coefficient) against a background of over 100,000 strictly neutral variants unlinked to the targets of selection. We filtered the dataset to retain only sites where the derived allele segregates at a minimum of two sampled time points, removing on average 21% of variants. As shown in the Supplementary Material, this filter does not affect inference accuracy, but it considerably reduces runtime.

#### Detection Power Evaluation

We evaluate the performance of `SweepLink` against three established single-locus tools: `ApproxWF` [Ferrer-Admetlla et al., 2016], `diplo-locus` [Cheng and Steinrücken, 2025], and `bmws` [Mathieson and Terhorst, 2022]. To ensure a fair comparison, the true simulated effective population size (*N* = 10,000) was explicitly provided as a fixed, known value to all methods. `SweepLink` was launched with 50 grid points for the numerical scheme of the diffusion equation and a discrete selection grid of 13 states for positive selection magnitudes from *s* = 0.0 to 0.06 with a step size of 0.005. This grid is extended in `SweepLink` to include corresponding negative selection values following Equation 1. The same complete grid is used for the maximum-likelihood approach in `diplo-locus`, which calculates the likelihood values across the grid and then interpolates them to find the off-grid maximum. We run `ApproxWF` and `bmws` with default parameters as recommended in their documentation. If not specified otherwise, MCMC in `ApproxWF` and `SweepLink` is run for 100,000 samples. For `ApproxWF`, we then discarded the first 10,000 samples as burning.

To compare the power of these tool to detect selection, we frame the problem as a binary classification task: how well does a tool distinguish neutral loci from those under directional selection? We extract different metrics for this classification task: For `diplo-locus`, we used the *p*-value derived from its likelihood ratio test [Cheng and Steinrücken, 2025]. For `bmws`, we follow the statistical framework described in its original publication Mathieson and Terhorst [2022]. Specifically, we aggregate single-locus selection estimations within sliding windows of 20 SNPs, approximate a gamma distribution, and derive a *p*-value by testing whether the window estimation deviates from this distribution. For `SweepLink` and `ApproxWF`, finally, we used MCMC-based Bayesian inference to yield posterior distributions for the selection coefficient. In its original implementation, `ApproxWF` considers loci to be under selection if the 95% credible interval of the posterior distribution excludes zero [Ferrer-Admetlla et al., 2016]. However, to ensure a fair comparison, we use the same score from Equation 7 as in `SweepLink`, defined as *P* = max(*P* (*s >* 0), *P* (*s <* 0)). We note that this metric for `ApproxWF` and `SweepLink` has an upper bound defined by the number *W* of MCMC samples: 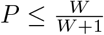.

To translate these continuous statistical scores into a binary classification, a formal discovery threshold must be applied. For example, a researcher might classify a locus as selected only if it achieves a *p*-value *<* 10^−7^ or a posterior probability *>* 0.99, treating all loci that fail to meet this threshold as neutral. An optimal threshold must maximize true positives (the correct identification of actual selective sweeps) while minimizing both false positives (the incorrect flagging of neutral loci) and false negatives (missed sweeps). To identify the optimal threshold and best performance for each tool, we evaluated the global classification accuracy across a continuous spectrum of all possible discovery thresholds. We quantified accuracy using the Matthews correlation coefficient (MCC) [Matthews, 1975] defined as

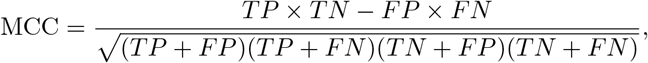

where *TP* represents the number of correctly identified loci under selection (True Positives), *TN* the number of correctly identified neutral loci (True Negatives), *FP* the number of neutral loci wrongly classified as selected (False Positives), and *FN* the number of selected loci wrongly classified as neutral (False Negatives). The MCC yields a value between −1 and 1, where 1 indicates perfect classification, 0 indicates performance no better than random guessing, and −1 indicates total disagreement between predictions and the true labels. We chose the MCC as an improvement over the classic *F*_1_ score, which can be misleading when applied to datasets with imbalanced classes [Chicco and Jurman, 2020], such as our simulated dataset containing roughly 80,000 neutral variants but only 500 selective sweeps.

Due to hitch-hiking, loci linked to loci under selection are not evolving neutrally and may not serve a neutral controls. We therefore evaluate detection performance in windows and classify a window as positive if at least one locus within it is significant given current threshold. To ensure independence, we split each chromosome into *k* = 21 windows (∼ 9,500 bp apart), yet only evaluate every second window. Of those, the middle window of a selected chromosome – the one containing the locus simulated to be under selection – is scored as a true positive (TP) if positive and a false negative (FN) otherwise. Every other window (eleven on any neutral chromosomes) are scored as a false positive (FP) if positive and a true negative (TN) otherwise.

We chose the method-specific thresholds such that all methods are aligned to an identical window-based False Positive Rate (FPR) of 0.3%, corresponding to the baseline error rate achieved by `SweepLink` at its optimal threshold. When comparing the accuracy in inferring selection strength, we compare the mean and variance of the estimated selection coefficients (*ŝ*) at the true locus, but excluded loci misclassified as neutral (false negatives).

### 2.8 SweepLink Robustness and Parameter Sensitivity

In a separate experimental setup, we evaluated the robustness and parameter sensitivity of the `SweepLink` framework under diverse evolutionary and sampling scenarios. We used data simulated by SLiM with a configuration described in previous section as a default configuration, but limited the experiments to three selection coefficients: weak *s* = 0.01, moderate *s* = 0.01 and strong *s* = 0.05. For these experiments, we thus simulated a total six chromosomes of 0.1 Mbp for each of the 100 replicates: one chromosome per included selection coefficient and three completely neutral chromosomes. To assess the impact of individual parameters on the accuracy of `SweepLink`, we then varied one parameter at a time, while keeping all other parameters constant at their default values. This way we investigated how the performance of `SweepLink` was affected by i) the sample size, ii) the sequence length, iii) the number of time points, iv) the starting allele frequency of the selective sweep, v) the recombination rate, and vi) the grid size in the numerical scheme for the diffusion equation. We further tested the effect of pooling samples from multiple generations into discrete time points, for which we simulated two diploid individuals for every of the 160 generations, and then compared the inference power using the correct sampling times against those when artificially pooling samples across generations into fewer effective time points. See the Supplementary Materials for further details and specific values tested.

### 2.9 Empirical Data

We applied `SweepLink` to the British ancient-DNA time series dataset that was previously analyzed using `bmws` in Mathieson and Terhorst [2022] and constructed from several resources [Mallick et al., 2024, version 44.3][Martiniano et al., 2016, Schiffels et al., 2016, Olalde et al., 2018, Brace et al., 2019, Margaryan et al., 2020, Patterson et al., 2022, Consortium et al., 2015]. This dataset contains 627 pseudo-haploid samples genotyped on 1240k SNPs, of which 535 are ancient. We converted sample dates to generations assuming 29 years per generation and binned the samples into 5-generation intervals, resulting in 26 time points spanning 0 − 150 generations before present (0 − 4,350 years). Sampling density is uneven across this range (Figure 2.9): the oldest sample dates to 4,480 BP and the youngest ancient sample to 930 BP, nearly half of all ancient chromosomes fall within ∼2,000 − 2,300 years BP. Starting from the original 1,150,639 autosomal SNPs, we remove 151 sites with no called genotypes and 144,019 sites that were monomorphic across the dataset.

We further removed sites with low-quality data: 437 sites at which more than 45% of the present-day samples were not called and an additional 149,822 sites at which more than 80% of the ancient samples were not called. Finally, to make our inference faster without losing valuable data, we remove 39,296 sites whose derived allele segregated at only one time point. We do not use MAF filtering as in the original study [Mathieson and Terhorst, 2022] to avoid bias in population size estimation. This leaves 816,914 SNPs genome-wide (71% of the original set).

We split the dataset by chromosome and used `SweepLink` to infer a selection coefficient at every locus together with an effective population size for each chromosome. A locus is called as potentially causal for selection when all 100,000 MCMC iterations agree on the sign of selection coefficient. At this chain length the criterion corresponds to a posterior probability of at least 0.99999, and in our simulations this maximum-stringency threshold gave `SweepLink` its best detection performance (Section 3.1). We then identify and report the selected region as the maximal run of consecutive SNPs at which the posterior probability of neutrality is below 0.5, and regard a region as affected by selection when it contains at least one locus that was detected as potentially causal variant for selection using the criterion above.

Loci identified as potentially causal variants for selection are carried through a common annotation pipeline. Associations were retrieved from OpenGWAS [Elsworth et al., 2020], which aggregates the NHGRI-EBI GWAS Catalog [**?**], UK Biobank [Bycroft et al., 2018] analyses with the MRC-IEU [Elsworth et al., 2019] and Neale laboratory pipelines [Neale Lab, 2018], and molecular-trait scans including eQTLGen whole-blood *cis*-eQTLs [**?**]. We queried every batch except FinnGen and BioBank Japan, whose populations differ too much from the British sample in linkage structure. Where associations were found, we tested whether the selection signal and the GWAS signal are consistent with a shared causal variant by approximate Bayes factor co-localisation, reporting the posterior probability of a common causal variant (PP_4_) [Giambartolomei et al., 2014]. We apply co-localisation analysis using a 1 Mb window centred on the target variant (*±* 500 kb) and the prior probabilities *p*_1_ = *p*_2_ = 10^−4^ and *p*_12_ = 10^−5^.

Detected variants were additionally annotated with the Ensembl Variant Effect Predictor [McLaren et al., 2016] to establish their functional consequence and their overlap with annotated regulatory features. For each region identified to be under selection, we plotted the inferred selection signal alongside the 1000 Genomes Phase 3 recombination map [Consortium et al., 2015, Howie et al., 2009] to check whether peaks coincided with regions of reduced recombination. For the lactase LCT region on chromosome 2, we additionally show the surrounding linkage disequilibrium block using the European block boundaries identified by LDetect Berisa and Pickrell [2015].

## 3 Results

### 3.1 Strong posterior support eliminates the need for threshold tuning

The window-based MCC values across the discovery thresholds for each tool are presented in Figure 3. `SweepLink` shows the highest peak performance (MCC = 0.87), ahead of `ApproxWF` (0.85), `diplo-locus` (0.86), and `bmws` (0.77). Each tool reaches its optimum at a threshold specific to its own score scale, but they differ in the shape of their threshold dependence and in how wide a range of thresholds sustains near-optimal performance. Since the optimal threshold is unknown in real empirical applications, the used threshold is unlikely optimal if the plateau is narrow. The region retaining performance within 0.01 MCC of the peak spans 1.6 orders of magnitude for `SweepLink` and 1.2 for `diplo-locus`, but only 0.7 for `ApproxWF` and 0.2 for `bmws`, after which performance declines steadily. At the high stringencies typically required for genome-wide scans (∼ 10^−7^), `diplo-locus` falls from its optimum of 0.86 to 0.78, and `bmws` from 0.77 to 0.07. `ApproxWF` reports a posterior probability rather than a *p*-value, so it is not subject to the same correction, but it is no less threshold-dependent: its optimum lies at a very lenient 1 − *P* ≈ 0.3, and demanding higher confidence costs it 0.09 MCC by 1 − *P* = 10^−4^.

**Figure 2.**
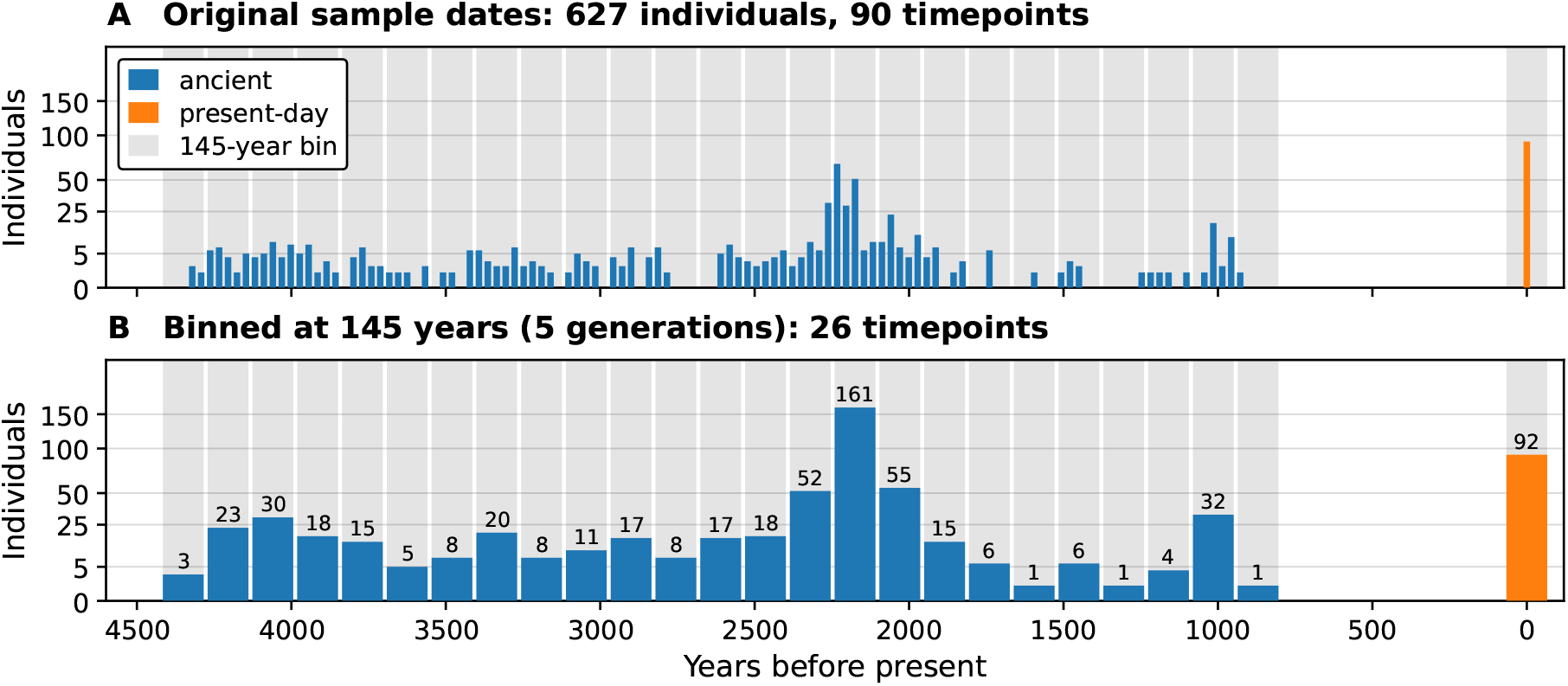
Temporal sampling of the British ancient-DNA dataset, before and after binning. **A** The dataset assembled from Mathieson and Terhorst [2022] contains 627 pseudo-haploid individuals at 90 distinct time points. **B** After binning by 5 generations of 29 years, 26 time points of 145 years remain. Shaded bands mark the bins and bars are labelled with the number of individuals they contain.

**Figure 3.**
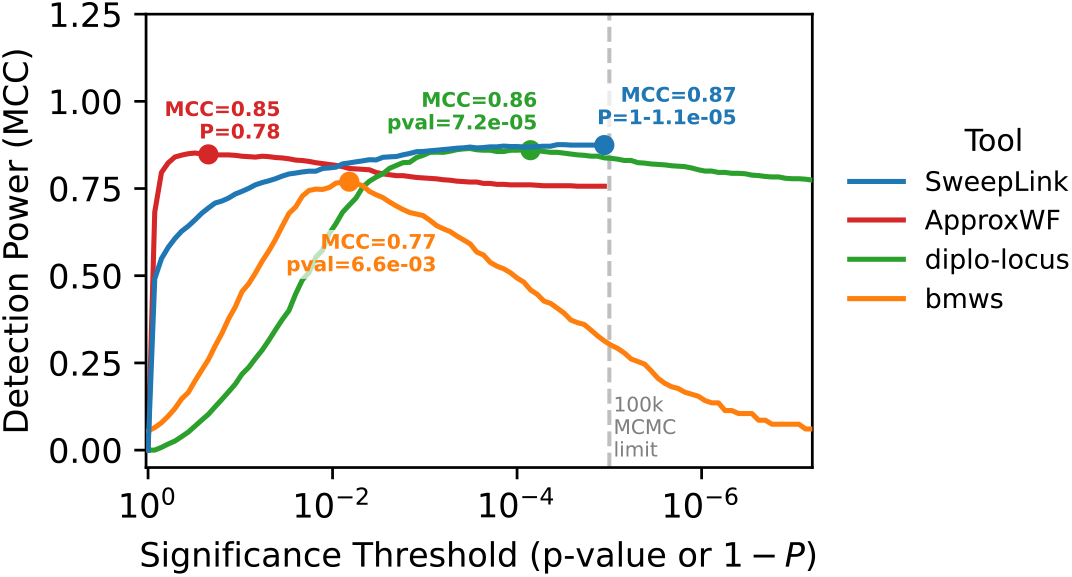
Power to detect selective sweeps across discovery thresholds. The x-axis displays native *p*-values (`diplo-locus`, `bmws`) or 1 − *P* (`SweepLink`, `ApproxWF`), where *P* = max(*P* (*s >* 0), *P* (*s <* 0)). Dots indicate peak MCC. Unlike standard methods, `SweepLink` receives peak accuracy at the maximally possible threshold (*P* = 1.0 − 10^−5^) in its default run.

`SweepLink` also reports posterior probabilities, but its optimum lies at the opposite end of the scale: it achieves the widest plateau of any tool tested and remains stable throughout the entire high-confidence domain (1 − *P <* 10^−4^): 0.87 at 10^−4^ and 0.88 at 10^−5^. Crucially, `SweepLink` reaches its optimal performance at the theoretical limit of maximum confidence (1 − *P* ∼ 10^−5^), which is set by the length of the MCMC chain: a posterior probability estimated from *W* samples cannot be resolved beyond 1*/*(1 + *W* ). Demanding the highest confidence the method can express is therefore also the best-performing choice, so no threshold tuning is required and the problem of an unknown optimum does not arise. The measured plateau width is consequently a lower bound rather than a property of the estimator: running an extended chain of 1,000,000 samples reproduces the same plateau and extends it by a further order of magnitude (Supplementary Figure **??**), confirming that the limit reflects the resolution of the estimator rather than its discriminative power.

As detailed in subsequent experiments, this robust plateau of `SweepLink` with peak at maximum threshold is consistently maintained across diverse evolutionary scenarios, bounded only by the fundamental limits of genetic linkage (Figure S6): at extremely high recombination rates (*r* ≥ 10^−6^ relative to a mutation rate of 10^−8^), the physical footprint of a sweep becomes too narrow to provide a meaningful regional context. In these isolated cases, the performance of `SweepLink` converges toward single-locus behaviour, exchanging its high-confidence plateau for a distinct performance peak at more lenient thresholds (Figure S6D).

### 3.2 Detection Sensitivity and Estimation Accuracy

Detection rates across four tools for five distinct categories of positive selection (*s* ∈ 0.01, 0.02, 0.03, 0.04, 0.05) are presented in Figure 4A, and the accuracy of the estimated selection coefficients (*ŝ*) for loci passing the matched window-based 0.3% FPR threshold are presented in Figure 4B. By *s* ≥ 0.04, all four tools reach 100% detection, and by *s* = 0.03 three of the four already do (`SweepLink`, `diplo-locus`, and `ApproxWF` at 100%, versus 91% for `bmws`). `SweepLink`’s advantage lies at weaker selection: at *s* = 0.02, it recovers 90% of sweeps, compared to 88% for `diplo-locus`, 80% for `ApproxWF`, and 31% for `bmws`, and at *s* = 0.01 these rates drop to 17%, 9%, 3%, and 0%, respectively.

**Figure 4.**
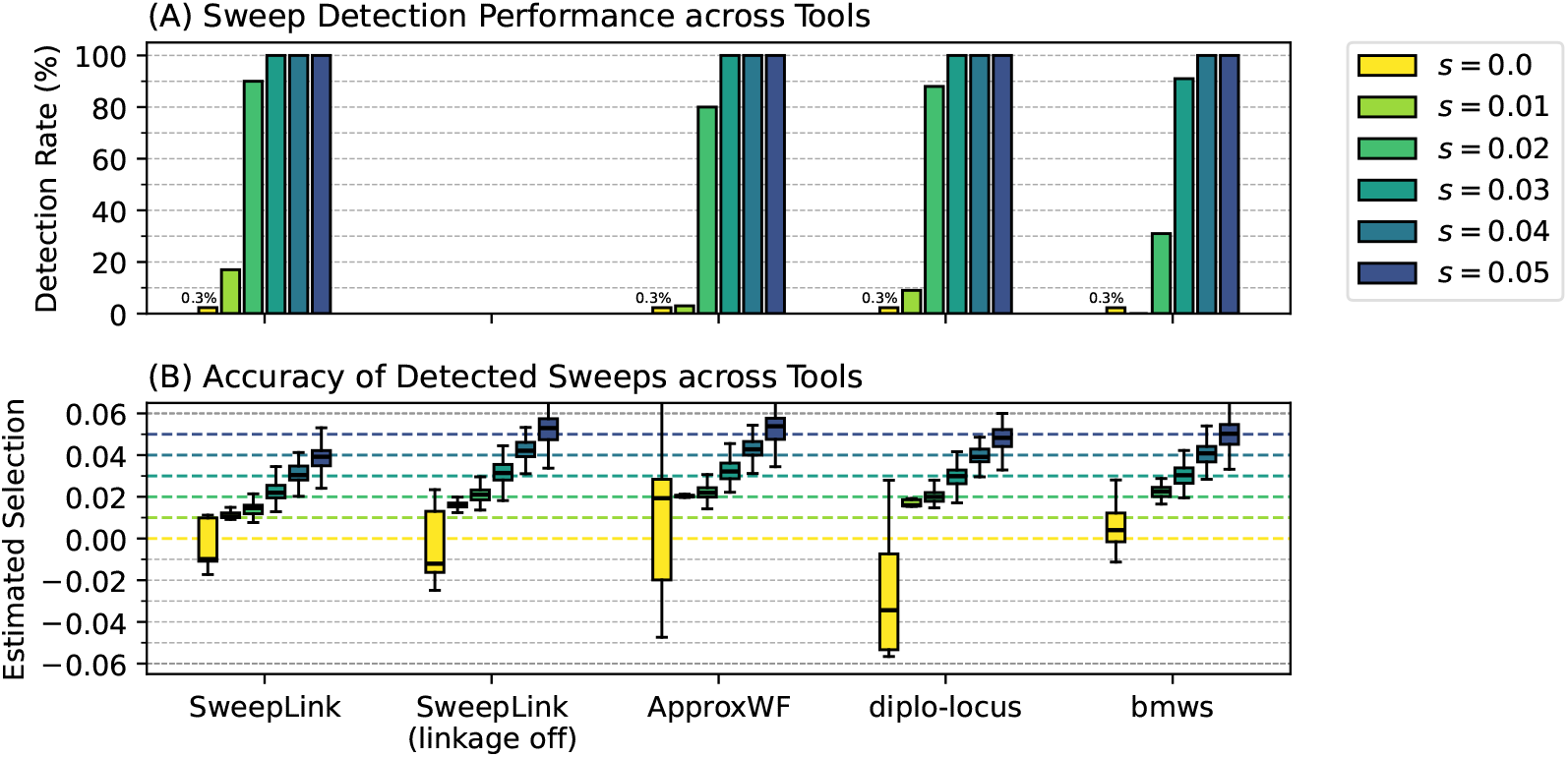
Performance and estimation accuracy for selection detection tools. **(A)** Sweep detection rates across selection coefficients evaluated at a matched window-based False Positive Rate (FPR) of 0.3% (the empirical FPR of `SweepLink` at 1 − *P* ∼ 10^−5^). `SweepLink` retains highest sensitivity for weak selection (*s* = 0.01). **(B)** Estimated selection coefficients (*ŝ*) for loci passing the 0.3% FPR threshold, including `SweepLink` estimates recomputed at the same detected loci with the spatial layer disabled (`SweepLink (linkage-off)`) to isolate linkage-pooling’s contribution to estimation bias. Coloured dashed lines indicate true values. Most existing tools overestimate selection magnitude for false positives (*s* = 0.0, yellow) and weak sweeps (*s* = 0.01, light green). In contrast, `SweepLink` tightly constrains neutral errors near zero and provides more accurate estimates for all selection coefficients, including weak selection (*s* = 0.01).

Yellow boxplots on Figure 4B show that `diplo-locus` and `ApproxWF` exhibit extreme variance when generating false positive calls, often incorrectly predicting high selection coefficients also for strictly neutral sites (*s* = 0.0). This behaviour is expected, as these tools analyse each locus independently and only detect selection when random genetic drift happens to mimic a strong selective sweep. The direction of this noise differs between the two: `ApproxWF` tends toward more positive estimates for its false positive calls, while `diplo-locus` produces high negative estimates – estimates that, because of occasional polarization errors flipping their sign, can become indistinguishable from a true signal at *s* = 0.04. At the same time, `bmws`’s window-based approach produces the estimations closest to zero for false positive calls, but only by sacrificing detection power, which is the lowest of all four tools. `SweepLink` avoids this trade-off entirely: it constrains its false-positive estimates far more tightly than `diplo-locus` or `ApproxWF`, and unlike `bmws`, does so without while retaining high detection power for real sweeps.

When comparing estimates for selected variants, competitive methods routinely overestimate weakly selected loci (*s* = 0.01), likely because weakly selected loci become significant only when random genetic drift magnifies the signal, producing a trajectory that mimics stronger selection (e.g., *s* ≈ 0.02). That effect vanishes for strongly selected loci, however, for which all methods have near 100% detection power and their median estiates of selection strength closely tracking simulated values.

In contrast, `SweepLink` provides highly accurate and unbiased estimates for weak selection (*s* = 0.01), which is both a result of its higher power to detect weakly selected loci, and because it pools information across neighbouring loci, which are unlikely to all show a consistently elevated selection signal due to genetic drift. This pooling of information, however, has the opposite effect at strongly selected loci (*s* ≥ 0.02), at which the neighbouring loci tend to have lower selection coefficients than the causal variant and pulling its estimate downwards. This effect is particularly pronounced at highly selected loci, for which the likelihood surface computed by `SweepLink` correctly peaks near the true value, yet remains relatively flat around this maximum, allowing it to be influences more heavily on the surrounding regional context. As a consequence, and while correctly ranking selected loci, `SweepLink` tends to systematically underestimate the selection coefficient and strongly selected sites.

To directly investigate the impact of accounting for the regional context, we re-estimated selection coefficients for all loci identified to be under selection using `SweepLink` with its spatial layer disabled, isolating each locus from its linked neighbours (population size again fixed to *N* = 10,000 and using a finer grid of *G* = 100 points due to the lower computational burden). The results, shown in Figure 4B (as `SweepLink (linkage off)`), confirm the trade-off: while ignoring information from neighbouring loci restores the accuracy of inferring strong selection coefficients, but results in an overestimation of weak selection coefficients.

### 3.3 Robustness

We evaluated how the accuracy of `SweepLink` in identifying selected loci is affected by sample size, number of time points, sequence length, the temporal binning strategy used to aggregate samples into a smaller number of discrete time points, initial frequency of the selected allele, recombination rate, and the numerical grid size used for the diffusion approximation. While we discuss these finding below, the MCC plots for each experiment are presented in Supplementary Figure S6.

Plots for detection power across selection coefficients, accuracy of selection estimations and posterior distributions for the population size are shown in Figure 5, Supplementary Figures S7–S12.

**Figure 5.**
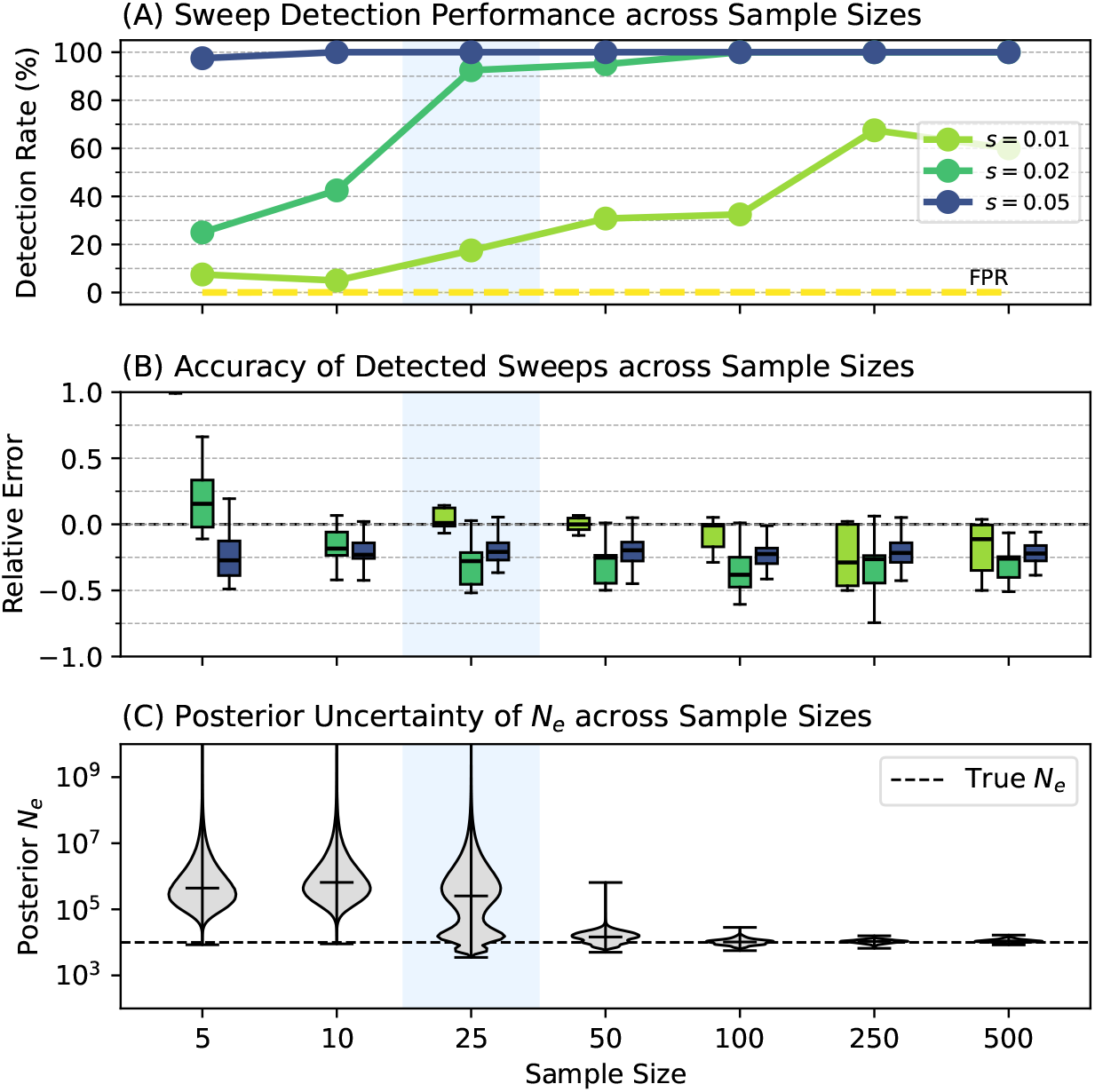
Needs caption.

The power to detect selection generally increases with larger sample size and more time points. In our default configuration with 25 diploid samples at 11 time point, `SweepLink` detected selection in 20% of cases for weak selection (*s* = 0.01), 81% for moderate selection (*s* = 0.02), and 100% for strong selection (*s* = 0.05), with an average locus-specific false positive rate of 0.09%. When repeating these experiments with 100 samples per time point, detection power for moderate selection reached 100%, and with 250 samples per time point, detection power for weak selection exceeded 50% (Figure 5A). Equally, increasing to 41 sampled time points increased detection power to 100% and above 40% for moderate and weak selection, respectively (Figure 5?).

Although neither sample size nor the number of time points had a consistent impact on the accuracy to infer selection coefficients (Figure 5B), they greatly improved the accuracy to infer the population size (Figure 5C and ?), for which the posterior distribution at low sample sizes and few time points was very broad and often included also unrealistically high values. Similarly to what was reported for `ApproxWF` [Ferrer-Admetlla et al., 2016], the likelihood surface computed by `SweepLink` from small sample sizes is very flat for large population sizes (Supplementary Figure S4), as the sampling variance is very large compared to the variance as a result of genetic drift, yet its peak is very close to the correct value of *N* = 10,000.

In line with previous findings [Ferrer-Admetlla et al., 2016], the initial allele frequency of the derived allele strongly affected detection power (Figure S11A), with intermediate frequencies yielding highest power because low frequency alleles are often lost by drift and high frequency alleles showing too small a frequency increase to be confidently distinguished from neutral alleles. As expected from `SweepLink`’s use of linkage information, selection was inferred more accurately in case of low recombination rates, and approached that of single-locus tools when recombination rates were high (Figure S10A). Chromosome length (i.e. the number of markers) did not affect selection-inference accuracy, but longer sequences yielded more concentrated population size posteriors (Supplementary Figure X).

Regarding technical aspects, we found `SweepLink` shows minimal sensitivity to the temporal binning of samples across multiple generations into one time point: detection power and population size accuracy remained stable across many bin sizes, though confidence intervals for population size widen as bins grew coarser, and at the coarsest binning tested (bin size 80, yielding only two bins and three time points) `SweepLink` consistently failed to recover the correct population size. While binning may be necessary due to imprecise dating, it may also result in fast calculations, and these results suggest that this is a fine strategy as long as bins are limited to a few generations. Finally, increasing the resolution of the numerical grid in the Wright-Fisher diffusion scheme generally improved `SweepLink`’s power to detect selection, but we found that as of a grid size of 50, the improvement was minimal and unlikely worth the additional computational costs.

### 3.4 Selection scan of the British ancient-DNA time series

As an illustration, we used `SweepLink` to jointly infer population size and per-locus selection coefficients in a human dataset of pseudo-haploid genotypes at 816,914 loci for 627 individuals sampled across the last four millennia on the British isles that was previously analysed with `bmws` [Mathieson and Terhorst, 2022]. Per chromosome effective population sizes estimates ranged from *N* = 6,023 to 14,148 across autosomes, consistent with published estimates for European populations for this period. The scan identified NN peaks containing at leas one locus with 1 − *P <* 10^−5^ 6, of which NN reached co-localisation support at PP_4_ *>* 0.8 (Table 1); the remainder are reported in Table 2.

**Table 1.**
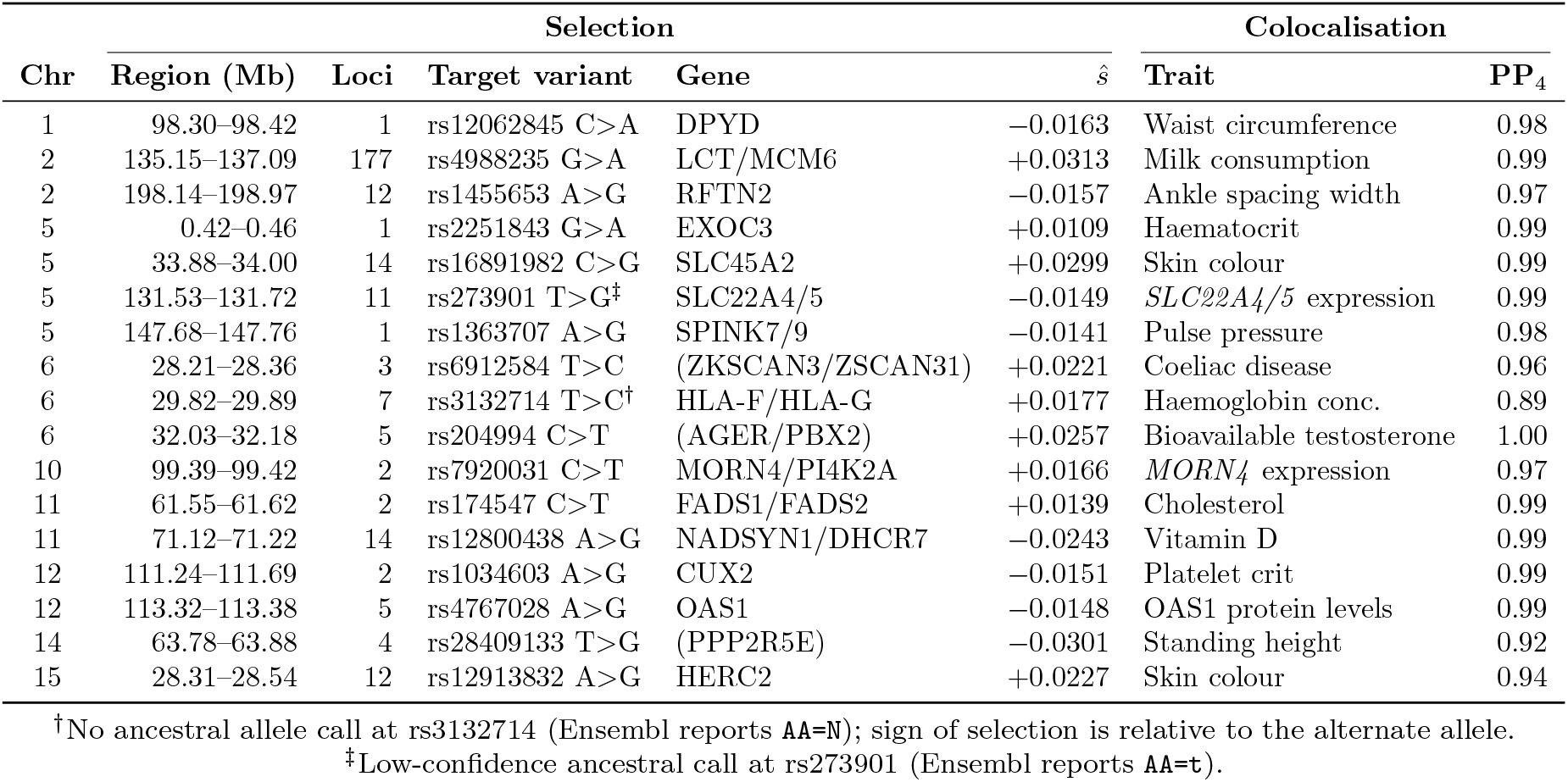
Regions under directed selection with co-localising trait associations. Selection coefficients (*ŝ*) refer to the derived allele, polarised against the Ensembl GRCh37 ancestral reconstruction; variants are written as ancestral*>*derived. “Loci” is the number of variants in the region at which all 10^5^ MCMC iterations agreed on the sign of *s*. Genes are the annotation at the region’s position; parentheses mark a target variant lying outside the named gene. PP_4_ is the posterior probability that the selection and trait signals share a causal variant.

**Table 2.** Regions under selection without independent functional support. *Tested* indicates co-localisation was run against every trait associated with the lead variant and no shared causal variant was recovered; *no assoc*. that no association reached genome-wide significance to test. *Corrupted p* marks a region whose only associated study reports *p*-values that are not measurements. *n* is the number of variants in the region.

| Chr | Region (Mb) | *n* | Lead SNP | *s* | Freq | Status |
| --- | --- | --- | --- | --- | --- | --- |
| 2 | 37.94–38.01 | 50 | rs72791933 | −0.0099 | 0.32 → 0.00 | No assoc. |
| 2 | 213.76–213.82 | 36 | rs10498001 | +0.0141 | 0.12 → 0.20 | Tested, PP4 0.00 |
| 2 | 221.74–221.79 | 18 | rs830747 | −0.0126 | 0.22 → 0.76 | No assoc. |
| 3 | 30.71–30.73 | 14 | rs3773656 | +0.0193 | 0.24 → 0.96 | No assoc. |
| 3 | 155.39–155.41 | 12 | rs358899 | +0.0184 | 0.77 → 0.92 | Tested, PP4 0.00 |
| 3 | 159.60–159.61 | 6 | rs3773785 | +0.0175 | 0.14 → 0.32 | Tested, PP4 0.00 |
| 3 | 184.38–184.41 | 18 | rs7653781 | +0.0155 | 0.91 → 0.95 | Tested, PP4 0.00 |
| 4 | 3.57–3.65 | 17 | rs73081157 | −0.0289 | 0.51 → 0.00 | No assoc. |
| 4 | 190.64–190.69 | 4 | rs11132751 | −0.0229 | 0.73 → 0.03 | No assoc. |
| 5 | 58.50–58.55 | 25 | rs66803649 | +0.0164 | 0.33 → 0.99 | No assoc. |
| 5 | 163.50–163.53 | 10 | rs10462943 | +0.0165 | 0.29 → 0.56 | Corrupted *p* |
| 5 | 179.22–179.37 | 45 | rs27471 | +0.0141 | 0.57 → 0.71 | No assoc. |
| 6 | 57.45–57.47 | 3 | rs9396343* | +0.0253 | 0.35 → 0.78 | No assoc. |
| 10 | 16.23–16.27 | 13 | rs4748263 | +0.0147 | 0.24 → 0.56 | No assoc. |
| 10 | 67.78–67.79 | 13 | rs6480128 | −0.0102 | 0.78 → 0.24 | No assoc. |
| 10 | 100.93–100.96 | 5 | rs6584238 | +0.0159 | 0.18 → 0.63 | No assoc. |
| 10 | 113.30–113.42 | 58 | rs11593093 | +0.0126 | 0.24 → 0.52 | No assoc. |
| 10 | 133.96–134.04 | 23 | rs1038804 | +0.0137 | 0.20 → 0.55 | Tested, PP4 0.67 |
| 11 | 7.96–7.98 | 17 | rs907203 | +0.0159 | 0.05 → 0.51 | No assoc. |
| 11 | 9.84–9.88 | 17 | rs2403221 | +0.0201 | 0.44 → 0.79 | Tested, PP4 0.29 |
| 11 | 34.94–34.95 | 26 | rs2956103 | −0.0105 | 0.71 → 0.21 | No assoc. |
| 11 | 105.93–105.95 | 3 | rs10895900 | +0.0234 | 0.20 → 0.75 | No assoc. |
| 12 | 78.30–78.35 | 20 | rs17817928 | −0.0116 | 0.29 → 0.10 | No assoc. |
| 12 | 93.06–93.10 | 14 | rs937754 | −0.0131 | 0.70 → 0.92 | No assoc. |
| 14 | 99.75–99.76 | 8 | rs5013941 | +0.0159 | 0.73 → 0.08 | No assoc. |
| 16 | 81.62 | 1 | rs4503787 | +0.0336 | 0.06 → 0.58 | Tested, PP4 0.50 |

\*Low-confidence ancestral call at rs9396343 (Ensembl reports AA=c); the magnitude of *s* is unaffected, the sign is not.

The scan recovered canonical targets of recent selection in Europeans. In line with previous findings [see Burger et al., 2020], lactase persistence at *LCT* /*MCM6* was the strongest signal in the genome, with 177 loci reaching the detection criterion across 1.9 Mb and a colocalisation with milk consumption at PP_4_ = 0.99. The scan further identified well known targets of selection in Europeans [], including the two major pigmentation loci *SLC45A2* (*ŝ* = +0.0310, skin colour, PP_4_ = 0.99) and *HERC2 ŝ* = +0.0227, skin colour, PP_4_ = 0.94), as well as the fatty acid desaturase cluster *FADS1* /*FADS2* (cholesterol, PP_4_ = 0.99), the vitamin D pathway at *NADSYN1* /*DHCR7* (PP_4_ = 0.99), as well as the antiviral *OAS1* locus, whose lead variant is a splice-acceptor change co-localising with OAS1 protein levels at PP_4_ = 0.99. The extended MHC on chromosome 6 contributes three further regions, co-localising respectively with coeliac disease (PP_4_ = 0.96), haemoglobin concentration (PP_4_ = 0.89) and bioavailable testosterone (PP_4_ = 1.00).

To explore the linkage structure of identified regions, we examined their local selection profiles and GWAS association tracks alongside population-matched recombination map from the 1000 Genomes Project (GBR population) [Consortium et al., 2015]. As illustrated in Figures **??**-**??**, this landscape analysis revealed a consistent pattern: the majority of our identified selection peaks are situated within recombination coldspots. Additionally, the selection footprint inferred at the *LCT* locus—a well-known target of selection for lactase persistence—aligns almost perfectly with the European LD block defined by LDetect [Berisa and Pickrell, 2015] (Figure 7).

**Figure 6.**
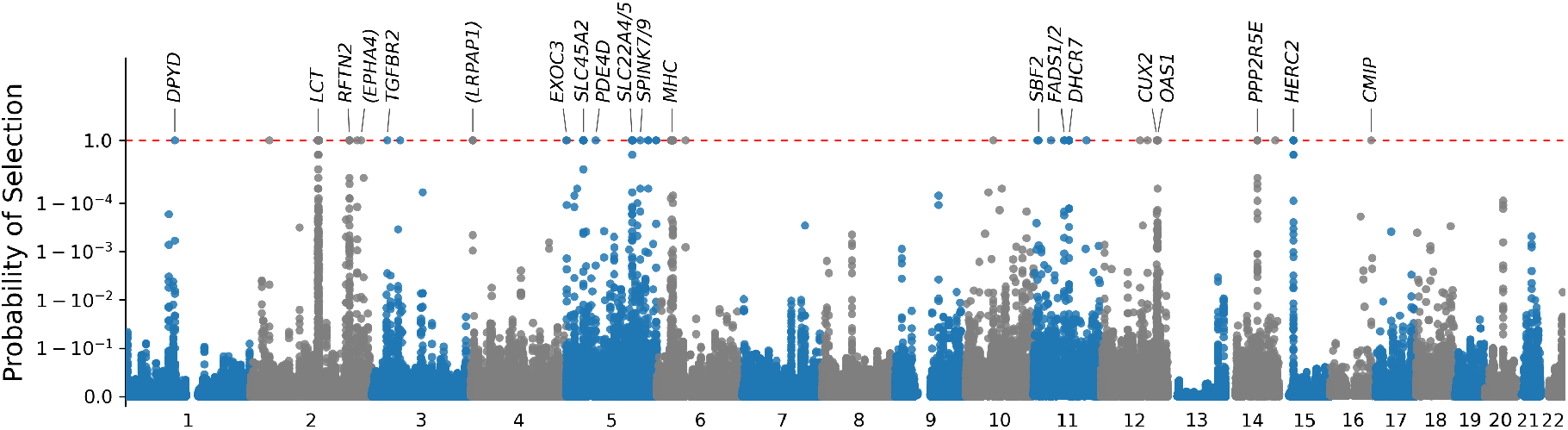
Genome-wide scan for signals of directed selection in Britain. The y-axis reflects the posterior one-sided probability of selection (*P* = max(*P* (*s >* 0), *P* (*s <* 0))), with the red dashed line representing the peak model confidence threshold (1 − *P* ∼ 10^−5^). Top-scoring variants within notable genomic regions are annotated with their respective candidate genes. Other prominent peaks with intergenic variants remain unannotated due to a lack of identifiable candidate genes or established GWAS associations at those loci.

**Figure 7.**
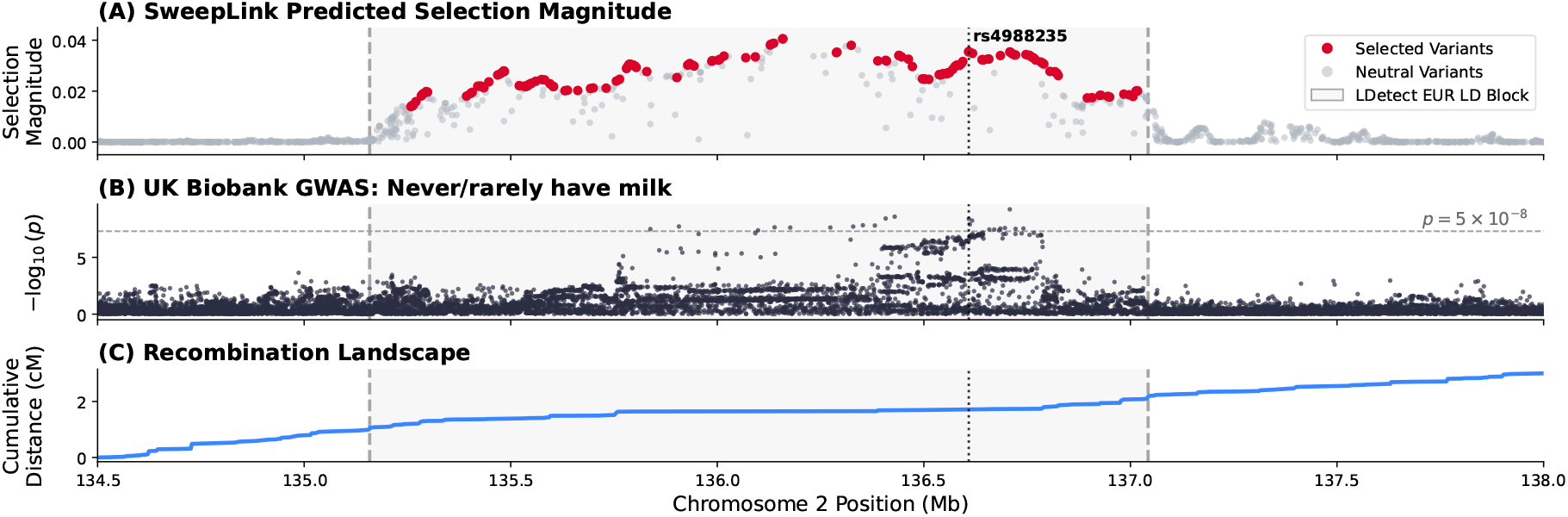
Genomic landscape of the selection signal at the *LCT* locus on chromosome 2. (A) Predicted selection magnitudes generated by `SweepLink`. Red points indicate variants classified as target variants under selection (*P* = 1.0), while gray points represent neutral variants. The known causal variant for lactase persistence, rs4988235, is marked by a vertical dotted line. The shaded background bounded by dashed vertical lines represents the European linkage disequilibrium (LD) block derived from the LDetect database. (B) Local Manhattan plot of GWAS summary statistics from the UK Biobank (Neale lab) for the trait “Milk type used: Never/rarely have milk” (OpenGWAS ID: ukb-d-1418 6). This trait colocalizes with predicted selection signal (colocalization probability = 0.99). The horizontal dashed line denotes the standard genome-wide significance threshold (*p* = 5 *×* 10^−8^). (C) Cumulative recombination landscape (cM) based on the 1000 Genomes Project genetic map for the GBR population.

We compared the results of `SweepLink` to a previous selection scan conducted on the same ancient British dataset [Mathieson and Terhorst, 2022]. That study reported seven genome-wide significant regions at their primary cutoff (*P <* 10^−7^), plus two further loci discussed separately: a locus at *OAS1* on chromosome 12, real but reaching significance only under a more lenient, properly Bonferroni-corrected threshold (*P* = 7.3 *×* 10^−7^), and a ninth signal on chromosome 4 near *LINC00955* that the authors themselves attribute to a likely genotyping artifact. `SweepLink` recovers seven of these eight real regions in some form, and correctly does not replicate the one they flagged as artifactual. Four are exact matches at the reported target variants: *LCT, SLC45A2, DHCR7*, and *HERC2* ; a fifth, *OAS1*, is likewise directly recovered. Within the extended MHC, two of our three detected peaks fall at essentially the same coordinates as two of the three HLA regions reported by Mathieson and Terhorst [2022]. The third of our MHC peaks (near *HLA-F* /*HLA-G* ) instead marks a different signal within the same broader region, and we do not recover their remaining HLA region. `SweepLink` additionally recovers the fatty acid desaturase cluster *FADS1* /*FADS2*, a locus that Mathieson and Terhorst [2022] themselves noted as suggestive but sub-threshold in their own scan (*P* = 1.5 *×* 10^−5^, against their 10^−7^ genome-wide cutoff). Beyond these previously reported loci, `SweepLink` identifies eight further candidate regions with independent co-localizing trait support that were not reported in the previous scan: *DPYD, RFTN2, EXOC3, SLC22A4* /*SLC22A5, SPINK7* /*SPINK9, MORN4* /*PI4K2A, CUX2*, and *PPP2R5E* (Table 1).

## 4 Discussion

Here we present `SweepLink`, the first method that jointly infers demography and linked selection from genome-wide time-series data. Due to generally low recombination rates, allele frequency trajectories of neighbouring loci are correlated [Hill and Robertson, 1968], particularly at selected sites, a phenomenon known as hitch-hiking [Barton, 2000]. Modelling this linkage structure is challenging, however, and thus often ignored [Ferrer-Admetlla et al., 2016, Cheng and Steinrücken, 2025, Foll et al., 2015, Mathieson and Terhorst, 2022], inferred for only pairs of loci [He et al., 2020], or requires extensive computational resources [Terhorst et al., 2015, Vaughn and Nielsen, 2024, Whitehouse and Schrider, 2023]. To address these challenges, we here introduce `SweepLink`, a novel approach, which does not attempt at modelling the correlation between linked explicitly, but at capturing the effect of hitch-hiking indirectly through the correlation of effective selection coefficients across the genome. This allows `SweepLink` to improve the detection power by both pooling evidence of selection across loci and down-weighting false signals caused by drift at isolated loci, while remaining computationally tractable also at the scale of genome-wide data.

Our simulation experiments demonstrated that `SweepLink` is more sensitive to weak and moderate selection than other tools, while its detection power for strong selection is the same. When `SweepLink` is able to capture the correlation between loci, it produces highly polarized posteriors that lead to the best performance at high stringency. Compared to other existing methods, this provides a practical advantage: other tools require an empirical threshold to separate neutral loci from selected ones and are highly dependent on this choice, whereas `SweepLink` provides its best estimates at its highest stringency level, thereby eliminating a dependence on threshold choice. In the absence of linkage, fo instance due to high recombination rates, `SweepLink` converges to the behaviour of single-locus tools and again requires threshold selection.

Our experiments on simulated data demonstrated the robustness of `SweepLink`’s performance across diverse evolutionary scenarios. Its accuracy and detection power grow with sample size, number of time points, and number of loci, while the temporal binning strategy has no significant effect on performance. Finally, as our recombination rate experiments show, `SweepLink`’s advantage over single-locus methods narrows once the recombination rate exceeds this range: as linkage disequilibrium decays too quickly to leave a detectable physical footprint around the causal variant, `SweepLink` converges to single-locus behaviour and its advantage over existing tools disappears.

The cost of the linkage correlation captured by `SweepLink` is reflected in the accuracy of its selection estimates: they are mostly underestimated for strong selection (*s* ≥ 0.02) because neighbouring loci tend to pull down the estimates at the causal locus. As we showed, this could potentially be corrected with a two-step approach: an initial run will all features enabled to detect loci under selection and infer demographic parameters with high detection power and low false-positive estimates. And an additional run with a finer grid and with the spatial (linkage) layer disabled to refine the estimates of selection coefficients on the subset of detected loci, while using the inferred population size.

Several limitations of the current model are worth noting. The spatial layer captures correlations only between neighboring loci through a first-order Markov structure, rather than the full joint co-variance across many linked loci. This trade-off is what makes genome-wide inference computationally tractable, but it also means `SweepLink` cannot represent longer-range linkage patterns that a richer, higher-order model might capture. However, since `SweepLink` does not directly model linkage but the correlation in selection coefficients, long-range effects are not broken by low-frequency variants, as their trajectories are usually compatible with any selection strength. `SweepLink` additionally assumes that both selection coefficients and demographic parameters (population sizes and migration rates) remained constant over the time period with samples. However, this could easily be relaxed under the diffusion framework implemented by implementing step changes at specific times.

Beyond demonstrating genome-scale scalability, the application to the British time series data on humans yielded substantive results in its own right. Our replicated signals and our novel candidates diverge in a revealing way. The loci `SweepLink` confirms (*LCT, SLC45A2, HERC2, DHCR7, OAS1*, and the matched HLA regions) all fit within the vitamin D/calcium selective-pressure hypothesis that Mathieson and Terhorst [2022] propose to explain nearly all of their British signals: lighter pigmentation and lactase persistence both increase vitamin D availability, *DHCR7* acts directly in its metabolism, and the HLA associations relate to celiac disease, itself a risk factor for calcium and vitamin D malabsorption. Our eight novel candidates, by contrast, co-localize with a more heterogeneous set of traits (including waist circumference, haematocrit, pulse pressure, and standing height) that do not obviously fit this narrative. This could indicate additional, previously uncaptured selective pressures acting on Bronze Age Britain. Equally, without a comparably coherent biological story to anchor them, we cannot rule out that some of these novel signals are false positives, and they should be treated as candidates for follow-up rather than as established findings.

`SweepLink`’s central practical advantage is also visible in how this scan was conducted. Mathieson and Terhorst [2022] chose a *P <* 10^−7^ significance cutoff that they describe as conservative for their number of tests, and validated post hoc using a permutation test on randomized sample dates. Our scan required no equivalent step: we simply applied `SweepLink` at its maximum-stringency posterior, the same default recommended by our simulation results, with no threshold tuning or post-hoc justification needed.

Several caveats apply to this application. Our co-localization analysis relies on present-day UK Biobank GWAS summary statistics, but several unsupported loci (Table 2) carry derived alleles that had drifted to near-zero or zero frequency by the present day and are therefore essentially untestable in a present-day cohort, regardless of whether they were genuinely under selection. As already noted by Mathieson and Terhorst [2022], this data set is also restricted to the 1240k capture panel used to genotype the ancient samples, so denser marker sets, whether from imputation or shotgun sequencing, could sharpen the localization of these signals or surface functional annotations currently missed. A small number of our sign-of-selection calls also carry ancestral-allele uncertainty: two loci in Table 1 (rs3132714, rs273901) have missing or low-confidence ancestral allele calls, so the direction, though not the magnitude, of the inferred selection coefficient at these sites should be treated with caution. Finally, like Mathieson and Terhorst [2022], we assume population continuity in Britain over this period; although Bronze Age migration into Britain is documented [Patterson et al., 2022], Mathieson and Terhorst [2022] argue this involved genetically similar, geographically adjacent populations unlikely to materially affect the interpretation of selection signals, an argument that applies equally to our results since we analyze the same dataset.

