## Supplemental Materials for "SweepLink: Joint Inference of Demography and Linked Selection from Time-series Data"

### Supplementary Material for *SweepLink: Joint Inference of Demography and Linked Selection from Time-series Data*

Ekaterina Noskova et al.

September 2, 2026

**7 Results for British data** **20**

### 1 Temporal Layer

#### 1.1 Wright-Fisher Diffusion

Assume $P$ populations. Consider **a single locus** with two alleles $A$ and $a$, where $a$ is the ancestral allele and $A$ is the derived allele, formed by mutation. Mutation acts on individual alleles at rates $\mu_{a\to A}$, from the ancestral allele $a$ to the derived allele $A$, and $\mu_{A\to a}$ for the back mutation. Denote $X_p(t) \in [0,1]$, $p = 1,\ldots,P$ — the relative frequency of the derived allele $A$ in a population $p$ at generation $t$. These frequencies form the random vector $\boldsymbol{X}(t) = \{X_p(t)\}_{p=1}^{P}$. According to the Wright-Fisher model $\boldsymbol{X}(t)$ is a Markov chain with a transition matrix defined by considered evolutionary forces such as population size, migration rate and selection rate.

Assume that each population $p$, $p = 1, \dots, P$, has an effective population size equal to $2N_p$. Migration between populations is described by the rates $0 \le m_{pq} \le 1$, where $m_{pq}$ is the migration rate from population $q$ to population $p$, satisfying

$$\sum_{q=1}^{P} m_{pq} = 1, \quad m_{pp} = 1 - \sum_{q \neq p} m_{pq}.$$

Selection at the given locus within population $q$ is governed by the fitness coefficients:

$$w_{AA} = 1 + s_q, \quad w_{Aa} = 1 + hs_q, \quad w_{aa} = 1,$$

where $s_q$ is the selection coefficient and $h$ the dominance coefficient equal across all populations.

To specify the transition of the Markov chain in the Wright-Fisher model, we calculate $\psi_p$, the probability of drawing a derived allele $A$ for population $p$ in generation $t+1$. Given the frequencies $\boldsymbol{X}(t) = \boldsymbol{x}$ at generation $t$, this probability is obtained through a three-stage sampling scheme. In the first stage, we select a parental population $q$ with probability $m_{pq}$. In the second stage, we draw an allele from the chosen population $q$ with a probability weighted by the fitness coefficients. Combining these two steps, the mean frequency of $A$ in population $p$ after migration and selection is

$$\phi_p(\boldsymbol{x}) = \sum_{q=1}^{P} m_{pq} \, \frac{w_{AA} x_q^2 + w_{Aa} x_q (1 - x_q)}{w_{AA} x_q^2 + 2 w_{Aa} x_q (1 - x_q) + w_{aa} (1 - x_q)^2}. \tag{S1}$$

In the third stage, we allow the sampled allele to mutate, which yields the final sampling probability

$$\psi_p(\boldsymbol{x}) = \tilde{x}_p + \mu_{a \to A}(1 - \phi_p(\boldsymbol{x})) - \mu_{A \to a} \, \phi_p(\boldsymbol{x}). \tag{S2}$$

The random variable $X_p(t+1)$ is then obtained by binomial sampling of $2N_p$ alleles with success probability $\psi_p$, as is standard in the Wright-Fisher model:

$$X_p(t+1) \;\sim\; \frac{\text{Binomial}(2N_p,\ \psi_p(\boldsymbol{X}(t)))}{2N_p}. \tag{S3}$$

For large $N_p$, the random vector $\boldsymbol{X}(t)$, which forms a discrete-time Markov chain, can be modeled approximatively as a multivariate diffusion process on the open cube $\mathcal{Q} = (0,1)^P$. One can show (see e.g. Karlin and Taylor (1981, ch. 15.2)) that the infinitesimal drift vector $\boldsymbol{a}(\boldsymbol{x})$ of the diffusion process is given by

$$a_p(\boldsymbol{x}) = \psi_p(\boldsymbol{x}) - x_p \tag{S4}$$

and the infinitesimal covariance matrix $\mathbf{b}(\boldsymbol{x})$ by

$$b_{pq}(\boldsymbol{x}) = \frac{x_p(1 - x_p)}{2N_p} \delta_{pq}, \tag{S5}$$

where $\delta_{pq}$ is the Kronecker delta. The infinitesimal generator of this process is the second-order linear operator:

$$\mathcal{L}_x \; f(t, \boldsymbol{x}) = \frac{1}{2} \sum_{p=1}^{P} b_{pp}(\boldsymbol{x}) \frac{\partial^2}{\partial {x_p}^2} f(t, \boldsymbol{x}) + \sum_{p=1}^{P} a_p(\boldsymbol{x}) \frac{\partial}{\partial x_p} f(t, \boldsymbol{x}), \tag{S6}$$

acting on some function $f(t, \boldsymbol{x})$. Its *formal* adjoint operator, acting on a function $g(t, \boldsymbol{y})$, is given by (see Karlin and Taylor (1981, ch. 15.5))

$$\mathcal{L}_y^* \; g(t, \boldsymbol{y}) = \frac{1}{2} \sum_{p=1}^{P} \frac{\partial^2}{\partial {y_p}^2} \left[ b_{pp}(\boldsymbol{y}) g(t, \boldsymbol{y}) \right] - \sum_{p=1}^{P} \frac{\partial}{\partial y_p} \left[ a_p(\boldsymbol{y}) g(t, \boldsymbol{y}) \right].$$

Consider the *transition probability density* $p(t_x, \boldsymbol{x}; t_y, \boldsymbol{y})$ to go from state $\boldsymbol{x}$ at time $t_x$ to state $\boldsymbol{y}$ at time $t_y$. Assume that changing $p(t_x, \boldsymbol{x}; t_y, \boldsymbol{y})$ is a stationary process (time homogeneous), i.e. $p(t_x, \boldsymbol{x}; t_x + \tau, \boldsymbol{y}) = p(0, \boldsymbol{x}; \tau, \boldsymbol{y})$, $\forall t_x, \tau > 0$. We define:

$$p(\tau, \boldsymbol{x}, \boldsymbol{y}) := p(t_x, \boldsymbol{x}; t_y, \boldsymbol{y})$$

for the transition time $\tau = t_y - t_x$.

Transition probability density $p(\tau, \boldsymbol{x}, \boldsymbol{y})$ satisfies the Fokker-Planck equation or **Kolmogorov's forward equation** $(t_y > t_x)$:

$$\frac{\partial}{\partial \tau} p(\tau, \boldsymbol{x}, \boldsymbol{y}) = \mathcal{L}_y^* \; p(\tau, \boldsymbol{x}, \boldsymbol{y}), \\ \text{subject to initial condition: } p(\tau, \boldsymbol{x}, \boldsymbol{y}) = \delta(\boldsymbol{y} - \boldsymbol{x}), \tag{S7}$$

where $\delta$ is the Dirac delta function. We have to prevent a flow of probability across the border of $\mathcal{Q} = [0, 1]^P$. The *probability current* $\boldsymbol{J}$ is defined as (see Gardiner et al. (1985, ch. 5.2.1))

$$J_p(\tau, \boldsymbol{y}) = \frac{1}{2} \frac{\partial}{\partial y_p} \left[ b_{pp}(\boldsymbol{y}) p(\tau, \boldsymbol{x}, \boldsymbol{y}) \right] - a_p(\boldsymbol{y}) p(\tau, \boldsymbol{x}, \boldsymbol{y})$$

where for notational convenience we suppressed the dependence on $\boldsymbol{x}$. We will set the **reflecting boundary conditions**:

$$J_p(\tau, \boldsymbol{y}) = 0 \quad \text{at } \boldsymbol{y} : \; y_q \in \{0, 1\}, \; q = 1, \ldots, P.$$

These ensure that the boundary of $\mathcal{Q}$ is a reflecting barrier and no probability mass can leak out from the interior of $\mathcal{Q}$. More about boundaries is available in Gardiner et al. (1985, ch. 5.2.1).

In addition, the transition probability density $p(\tau, \boldsymbol{x}, \boldsymbol{y})$ satisfies **Kolmogorov's backward equation** with respect to the variable $x$:

$$\frac{\partial}{\partial \tau} p(\tau, \boldsymbol{x}, \boldsymbol{y}) = \mathcal{L}_x \; p(\tau, \boldsymbol{x}, \boldsymbol{y}), \\ \text{subject to initial condition: } p(\tau, \boldsymbol{x}, \boldsymbol{y}) = \delta(\boldsymbol{x} - \boldsymbol{y}). \tag{S8}$$

The boundary conditions for a **reflecting barrier** are of Neumann type (see Gardiner et al. (1985, ch. 5.2.1)):

$$\frac{\partial}{\partial x_p} p(\tau, \boldsymbol{x}, \boldsymbol{y}) = 0, \quad \text{at } \boldsymbol{x} : \; x_q \in \{0, 1\}, \; q = 1, \ldots, P.$$

It can be shown that $\mathcal{L}_x$ and $\mathcal{L}_y^\star$ with described reflecting boundary conditions are *truly adjoint* operators in $L^2([0, 1])$.

#### 1.2 Hidden Diffusion Model

We now imagine our diffusion process $\boldsymbol{X}(t)$ is a hidden process of the HMM. At given times $t_0 < t_2 < \ldots < t_K$ we have $n_{pk}$ **diploid** samples of our observed locus, where $p = 1, \ldots, P$ — population number and $k = 0, \ldots, K$ — index of corresponding time point.

Consider $c_{pk} \in [0, 2n_{pk}]$ — the absolute frequency of allele $A$ among samples of the population $p$ at a time point $t_k$. Each $c_{pk}$ is an independent binomial variable, conditional on the unobserved value of $X_p(t_k) = x_{pk}$. More precisely, the emission probabilities for the sampling process $\boldsymbol{C}(t_k)$ are:

$$\mathcal{B}(\mathbf{c}_k | \mathbf{x}_k) := \mathbb{P} \left[ \mathbf{C}(t_k) = \mathbf{c}_k | \mathbf{X}(t_k) = \mathbf{x}_k \right] = \prod_{p=1}^{P} \binom{2n_{pk}}{c_{pk}} x_{pk}^{c_{pk}} (1 - x_{pk})^{2n_{pk} - c_{pk}}. \tag{S9}$$

##### 1.2.1 Forward Recursion

Consider a time point $t$ between $t_k$ and $t_{k+1}$: $t_k \leq t < t_{k+1}$. Let $\tau = t - t_k$. Denote by $\alpha_k(\tau, \boldsymbol{x})$ the probability of observing the emission data $\mathbf{c}_{0:k}$ up to $t_k$ and to be in state $\boldsymbol{y}$ at time $t = t_k + \tau$ (Figure S1):

$$\alpha_k(\tau, \boldsymbol{y}) := \mathbb{P} \left[ \boldsymbol{X}(t_k + \tau) = \boldsymbol{y}, \boldsymbol{C}(t_0) = \mathbf{c}_0, \ldots, \boldsymbol{C}(t_k) = \mathbf{c}_k \right].$$

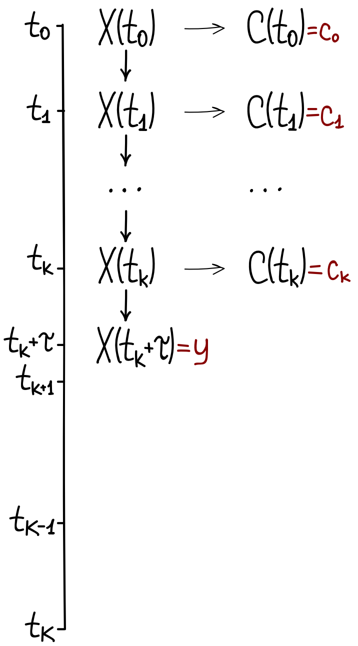

Figure S1: Schematic of the forward variable $\alpha_k(\tau, \boldsymbol{y})$. Time runs downward from $t_0$ to $t_K$. The hidden allele frequencies $\boldsymbol{X}(t)$ (centre) evolve through the Wright-Fisher diffusion and emit the observations $\boldsymbol{C}(t)$ of the allele counts (right). The quantities in dark red are those whose joint probability defines $\alpha_k(\tau, \boldsymbol{y})$: the past and present observations $\mathbf{c}_0, \ldots, \mathbf{c}_k$, together with the process state $\boldsymbol{y}$ at the intermediate time $t_k + \tau$. The future time points $t_{k+1}, \ldots, t_K$ are left blank, as their observations do not define $\alpha_k$.

The $\alpha_k$ satisfy the forward recursion

$$\alpha_k(\tau, \boldsymbol{y}) = \int\limits_{\mathcal{Q}} \alpha_{k-1}(\Delta t_{k-1}, \boldsymbol{x}) \mathcal{B}(\mathbf{c}_k|\boldsymbol{x}) p(\tau, \boldsymbol{x}, \boldsymbol{y}) \, \mathrm{d}\boldsymbol{x}, \tag{S10}$$

where $\Delta t_{k-1} = t_k - t_{k-1}$. From Equation S10 and Equation S7 we can derive that $\alpha_k(\tau, \boldsymbol{y})$ satisfies the forward Kolmogorov equation:

$$\frac{\partial}{\partial \tau} \alpha_k(\tau, \boldsymbol{y}) = \mathcal{L}_y^* \; \alpha_k(\tau, \boldsymbol{y}) \tag{S11}$$

$$\text{subject to initial condition: } \alpha_k(0, \boldsymbol{y}) = \alpha_{k-1}(\Delta t_{k-1}, \boldsymbol{y}) \mathcal{B}(\mathbf{c}_k|\boldsymbol{y}) \tag{S12}$$

and reflecting boundary conditions:

$$\frac{1}{2} \frac{\partial}{\partial y_p} \left[ b_{pp}(\boldsymbol{y}) \alpha_k(\tau, \boldsymbol{y}) \right] - a_p(\boldsymbol{y}) \alpha_k(\tau, \boldsymbol{y}) = 0 \quad \text{at } \boldsymbol{y} : \; y_q \in \{0, 1\}, \; q = 1, \ldots, P.$$

The initial condition for $\alpha_0(\tau, \boldsymbol{y})$ is special, namely

$$\alpha_0(0, \boldsymbol{y}) = \alpha(\boldsymbol{y}) \mathcal{B}(\mathbf{c}_0|\boldsymbol{y}) \tag{S13}$$

where $\alpha$ is a prior density on $\mathcal{Q}$.

That said, we can now, inductively and starting with $\alpha_0$, determine the $\alpha_k$ by numerically solving each PDE S10 on its time interval $0 \leq \tau \leq \Delta t_k$, using the previous solution for the new initial condition.

##### 1.2.2 Backward Recursion

Consider a time point $t$ between $t_k$ and $t_{k+1}$: $t_k < t \leq t_{k+1}$. This time, let $\tau = t_{k+1} - t$. Denote by $\beta_k(\tau, \boldsymbol{x})$ the probability of observing the future emission data $\mathbf{c}_{k+1:K}$ conditional to being in state $\boldsymbol{x}$ at time $t$ (Figure S2):

$$\beta_k(\tau, \boldsymbol{x}) := \mathbb{P}\left[\mathbf{C}(t_{k+1}) = \mathbf{c}_{k+1}, \ldots, \mathbf{C}(t_K) = \mathbf{c}_K \middle| \boldsymbol{X}(t_{k+1} - \tau) = \boldsymbol{x}\right].$$

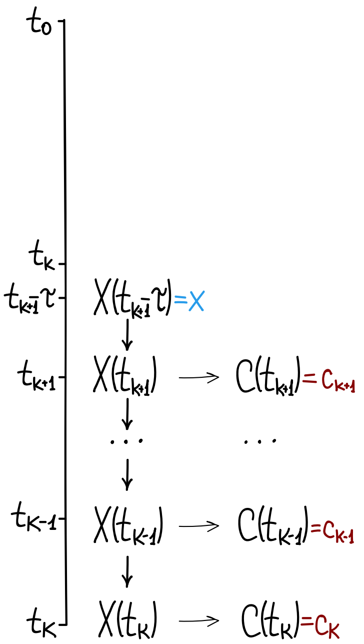

Figure S2: Schematic of the backward variable $\beta_k(\tau, \boldsymbol{x})$. Time again runs downward from $t_0$ to $t_K$. Conditional on the process occupying state $\boldsymbol{x}$ (blue) at the intermediate time $t_{k+1} - \tau$, the variable $\beta_k(\tau, \boldsymbol{x})$ gives the probability of the future observations $\mathbf{c}_{k+1}, \ldots, \mathbf{c}_K$ (red). The earlier time points are left blank, as the past observations do not define $\beta_k$. Together with Figure **??**, this illustrates the complementary roles of the forward and backward recursions.

The $\beta_k$ satisfy the backward recursion:

$$\beta_k(\tau, \boldsymbol{x}) = \int_{\mathcal{Q}} p(\tau, \boldsymbol{x}, \boldsymbol{y}) \mathcal{B}(\mathbf{c}_{k+1}|\boldsymbol{y}) \beta_{k+1}(\Delta t_{k+1}, \boldsymbol{y}) \, \mathrm{d}\boldsymbol{y}, \tag{S14}$$

where $\Delta t_{k+1} = t_{k+2} - t_{k+1}$. From Equation S14 and Equation S8 we can derive that $\beta_k(\tau, \boldsymbol{x})$ satisfies Kolmogorov's backward equation:

$$\frac{\partial}{\partial \tau} \beta_k(\tau, \boldsymbol{x}) = \mathcal{L}_x \; \beta_k(\tau, \boldsymbol{x}) \tag{S15}$$

$$\text{subject to initial condition: } \beta_k(0, \boldsymbol{x}) = \mathcal{B}(\mathbf{c}_{k+1}|\boldsymbol{x}) \beta_{k+1}(\Delta t_{k+1}, \boldsymbol{x}) \tag{S16}$$

and the (Neumann type) reflecting boundary conditions:

$$\frac{\partial}{\partial x_p} \beta_k(\tau, \boldsymbol{x}) = 0, \quad \text{at } \boldsymbol{x} : \; x_q \in \{0, 1\}, \; q = 1, \ldots, P.$$

The initial condition for $\beta_{K-1}(\tau, \boldsymbol{x})$ is

$$\beta_{K-1}(0, \boldsymbol{x}) = \mathcal{B}(\mathbf{c}_K|\boldsymbol{x}). \tag{S17}$$

Recursively and starting with $\beta_{K-1}$, we determine the $\beta_k$ by numerically solving each PDE S14 on its time interval $0 \leq \tau \leq \Delta t_k$, using the previous solution for the new initial condition.

##### 1.2.3 Likelihood

Denote population sizes, selection, migration rates as parameters $\theta$. Those parameters form an HMM transition and emission probabilities. Having either $\alpha_k$ or $\beta_k$ we can compute the probability of getting the observations $c_0, \ldots, c_K$ given parameters $\theta$.

Using forward recursion:

$$L(\theta \,|\, \mathbf{c}) = \mathbb{P}(C(t_0) = c_0, \ldots, C(t_K) = c_K \,\big|\, \theta) = \int\limits_{\mathcal{Q}} \alpha_K(0, \boldsymbol{y})\, d\boldsymbol{y} \tag{S18}$$

Using backward recursion:

$$L(\theta \,|\, \mathbf{c}) = \mathbb{P}(C(t_0) = c_0, \ldots, C(t_K) = c_K \,\big|\, \theta) = \int\limits_{\mathcal{Q}} \beta_0(\Delta t_0, \boldsymbol{x}) \mathcal{B}(\mathbf{c}_0 | \boldsymbol{x})\, d\boldsymbol{x} \tag{S19}$$

##### 1.2.4 Approach Summary

For clarity, we write down the procedure we follow to compute the likelihood $L(\theta \,|\, \mathbf{c}) = \mathbb{P}(C(t_0) = c_0, \ldots, C(t_K) = c_K \,\big|\, \theta)$ for given values of parameters $\theta = (N_p, s_p, h, m_{pq})_{p,q=1}^{P}$. It can be done using either forward or backward recursion.

Using forward:

1. Build infinitesimal mean $a_p(\boldsymbol{y})$ and variance $b_{pp}(\boldsymbol{y})$ according to the given $\theta$.
2. Set $k = 0$ and repeat until $k = K - 1$:
   (a) Construct the initial condition $\alpha_k(0, \boldsymbol{y})$ using Equation S13 if $k = 0$ or Equation S12 otherwise.
   (b) Obtain $\alpha_k(t_{k+1} - t_k, \boldsymbol{y})$ for $\boldsymbol{y} \in [0, 1]^P$ by numerically solving the KFE in Equation S11 forward in time for $\tau \in [0, t_{k+1} - t_k]$ with initial condition $\alpha_k(0, \boldsymbol{y})$ and reflecting boundary conditions.
   (c) Set $k = k + 1$.
3. Obtain $\alpha_K(0, \boldsymbol{y})$ using Equation S12.
4. Combine $\alpha_K(0, \boldsymbol{y})$ for Equation S18 to obtain $L(\theta \,|\, \mathbf{c})$.

Using backward:

1. Build infinitesimal mean $a_p(\boldsymbol{y})$ and variance $b_{pp}(\boldsymbol{y})$ according to the given $\theta$.
2. Set $k = K - 1$ and repeat until $k = 0$:
   (a) Construct the initial condition $\beta_k(0, \boldsymbol{x})$ using Equation S17 if $k = K - 1$ or Equation S16 otherwise.
   (b) Obtain $\beta_k(t_{k+1} - t_k, \boldsymbol{x})$ for $\boldsymbol{x} \in [0, 1]^P$ by numerically solving the KBE in Equation S15 forward in time for $\tau \in [0, t_{k+1} - t_k]$ with initial condition $\beta_k(0, \boldsymbol{x})$ and reflecting boundary conditions.
   (c) Set $k = k - 1$.
3. Combine $\beta_0(t_1 - t_0, \boldsymbol{x})$ for Equation S19 to obtain $L(\theta \,|\, \mathbf{c})$.

### 2 Overview of Numerical Methods

To compute the likelihood of the Hidden Markov Model, one can evaluate either the forward recursion (Eq. S18) or the backward recursion (Eq. S19). Initially, we implemented the Crank-Nicolson scheme for both the forward and backward Kolmogorov equations.

Although the Crank-Nicolson scheme is unconditionally stable, it often produces spurious oscillations when the drift term dominates the diffusion term or when sharp gradients occur (Morton and Mayers, 2005). This can result in non-physical negative probabilities, which break Hidden Markov Model likelihood calculations. To avoid this, we use the Chang-Cooper scheme (Chang and Cooper, 1970; Gutenkunst et al., 2009) for the forward pass, as it naturally conserves probability mass and strictly guarantees non-negative solutions.

We compute all final empirical results in this study using the forward recursion paired with the robust Chang-Cooper scheme. For methodological completeness, we provide the derivations for the Crank-Nicolson scheme for the backward recursion.

#### 2.1 Grid Discretization

Assume the time axis is divided into $M$ intervals, yielding $M+1$ grid points, and the frequency axis for a specific population is divided into $D+1$ intervals, yielding $D+2$ grid points and $D$ inner grid points. The grid points are denoted by:

$$\begin{aligned} \tau_n, &\quad \text{for } n = 0, 1, \ldots, M, \\ y_i, &\quad \text{for } i = 0, 1, \ldots, D+1. \end{aligned}$$

The grid spacing is given by:

$$\begin{aligned} \Delta\tau_n = \tau_n - \tau_{n-1}, &\quad \text{for } n = 1, \ldots, M, \\ \Delta_i = y_i - y_{i-1}, &\quad \text{for } i = 1, \ldots, D+1. \end{aligned}$$

While time steps $\Delta\tau_n$ are typically uniform, the spatial grid often benefits from clustering points near the boundaries 0 and 1 to properly capture the effects of genetic drift Gutenkunst et al. (2009); Fine and Steinrücken (2025). Therefore, we constructed and tested several symmetrical spatial grids that vary in their clustering density near the boundaries. The implemented grid mapping functions are (Figure S3):

- **Uniform grid.**

$$y_i = \xi_i = \frac{i}{D+1}.$$

- **Quadratic grid.** Adapted from other population genetics studies (Gutenkunst et al., 2009; Malaspinas et al., 2012; Ferrer-Admetlla et al., 2016), this grid is designed such that the spacing between adjacent points follows a quadratic density, clustering grid points near the boundaries. Compared to previous implementations, we construct a strictly symmetric and continuous grid. Starting with $y_0 = 0$:

$$y_i = y_{i-1} + x_i(1 - x_i), \quad \text{where} \quad x_i = \frac{i - 0.5}{D+1},$$

  the grid is subsequently scaled so that $y_{D+1} = 1$.

- **Chebyshev grid.** Based on the roots of Chebyshev polynomials, these nodes minimize Runge's phenomenon in numerical integration (Mathews et al., 2004). Their application for population genetic time-series models was recently shown in Fine and Steinrücken (2025). The ascending grid points on the interval $[0, 1]$ are given by:

$$y_i = \frac{1}{2}\left[1 - \cos\left(\pi\xi_i\right)\right]$$

- **Logistic grid.** Also referred to as an exponential grid (Malaspinas et al., 2012), it uses the following logistic function defined by a scale parameter $a > 0$:

$$y_i = \frac{1}{1 + e^{-a\cdot(\xi_i - 0.5)}},$$

  the grid is further scaled so that $y_0 = 0$ and $y_{D+1} = 1$.

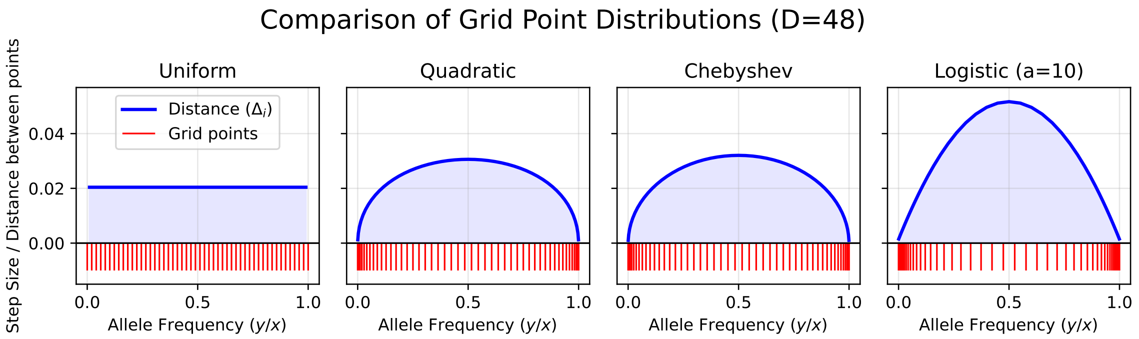

Figure S3: **Comparison of implemented spatial grid types for allele frequencies.** Four symmetric grid types are shown using $D = 48$ internal points (50 total grid nodes). The red vertical lines (bottom) indicate the exact locations of the grid nodes. The solid blue curves represent the step size ($\Delta_i$) between adjacent nodes. While the Uniform grid maintains a constant step size, the Quadratic, Chebyshev, and Logistic ($a = 10$) grids exhibit varying degrees of density clustering near the boundaries (0 and 1).

#### 2.2 1D Forward Equation (Chang-Cooper Scheme)

First, we derive numerical solution for Kolmogorov forward equation in case of one population. We later extend it to the multi-dimensional case.

For notational simplicity, we drop the population and time segment indices, focusing on the evaluation of the density function $\alpha(\tau, y)$ over time. We remind that we solve the following equation:

$$\frac{\partial}{\partial \tau}\alpha(\tau, y) = \frac{1}{2}\frac{\partial^2}{\partial y^2}\left[b(y)\alpha(\tau, y)\right] - \frac{\partial}{\partial y}\left[a(y)\alpha(\tau, y)\right],$$

$$\text{with a boundary condition: } \frac{1}{2}\frac{\partial}{\partial y}\left[b(y)\alpha(\tau, y)\right] - a(y)\alpha(\tau, y) = 0,$$

where function $a(y)$ and $b(y)$ are defined by Equation S4 and Equation S5 correspondingly.

The Chang-Cooper scheme is specifically engineered for Fokker-Planck equations, guaranteeing mass conservation and non-negative solutions (Chang and Cooper, 1970). To use the same notation as in the original paper, we rewrite the forward Kolmogorov equation in the following flux form:

$$\frac{\partial}{\partial \tau}\alpha(\tau, y) = \frac{\partial}{\partial y}\left[C(y)\frac{\partial}{\partial y}\alpha(\tau, y) + B(y)\alpha(\tau, y)\right] = \frac{\partial F}{\partial y},$$

where the substituted drift and diffusion functions are defined as:

$$B(y) = \frac{1}{2}\frac{\partial}{\partial y}[b(y)] - a(y),$$

$$C(y) = \frac{1}{2}b(y).$$

Having Equation S5 for $b(y)$, we evaluate the analytical derivative as:

$$\frac{\partial}{\partial y}[b(y)] = \frac{1 - 2y}{2N}$$

Regarding the boundary condition, we note that $b(y = 0)$ and $b(y = 1)$ evaluate to 0, while $a(y) \neq 0$ at the boundaries. Given our zero-flux reflecting boundary condition, this yields $\alpha(y = 0) = 0$ and $\alpha(y = 1) = 0$. The Chang-Cooper scheme is a finite-volume method and operates using numerical fluxes — the rate of probability mass flowing across the boundaries of each cell. By explicitly setting the exterior boundary fluxes $F_{1/2}$ and $F_{D+1/2}$ to 0, we can safely enforce the reflecting boundary conditions. This reduces our numerical system to a $D \times D$ matrix operating exclusively on the interior grid points.

Chang-Cooper scheme discretize the equation in the following way:

$$\frac{1}{\Delta\tau_n}\left(\alpha_j^{n+1}-\alpha_j^n\right)=\frac{1}{\Delta_j}\left(F_{j+1/2}-F_{j-1/2}\right), \tag{S20}$$

where $\alpha_j^n=\alpha(\tau^n,y_j)$ and flux $F_{j+1/2}$ is defined as:

$$\begin{aligned}F_{j+1/2}&=\left((1-\delta_j)B(y_{j+1/2})+\frac{1}{\Delta x}C(y_{j+1/2})\right)\alpha_{j+1}^{n+1}-\left(\frac{1}{\Delta x}C(y_{j+1/2})-\delta_j B(y_{j+1/2})\right)\alpha_j^{n+1}\\ F_{1/2}&=F_{D+1/2}=0.\end{aligned} \tag{S21}$$

This weighted average relies on a dynamic flux-weighting parameter $\delta_j\in[0,1]$. This parameter adaptively shifts the discrete approximation from a centered differencing scheme (when $\delta_j=0.5$) to an upwind scheme (as $\delta_j$ approaches 0 or 1). It is defined in such a way that the method yields stable, non-negative solutions, without restriction on $\Delta_j$. In the original paper Chang and Cooper (1970) this coefficient is defined as:

$$\begin{aligned}\delta_j^{n+1}&=\frac{1}{\omega_j}+\frac{1}{1-\exp(\omega_j)},\\ \omega_j&=\frac{B(y_{j+1/2})}{C(y_{j+1/2})}\Delta_{j+1}\end{aligned}$$

However, our empirical experiments demonstrated that evaluating $\omega$ as a ratio at the midpoint is unstable for the Wright-Fisher model. This occurs because the diffusion coefficient $C(y)$ goes to 0 at the boundaries 0 and 1. The stability can be improved by using non-uniform grids with points clustered at the boundaries. However, we adopted the structure-preserving Chang-Cooper (SP-CC) framework proposed by Pareschi and Zanella (2018) instead:

$$\omega_j=\int_{y_j}^{y_{j+1}}\frac{B(y)}{C(y)}\,dy=\int\frac{B_p(y)}{C_p(y)}\bigg|_{y_j}^{y_{j+1}} \tag{S22}$$

Substituting our specific drift and diffusion functions, this integral yields an exact, closed-form analytical solution. Reintroducing the population index $p$, the general antiderivative takes the form:

$$\begin{aligned}\int\frac{B_p(y)}{C_p(y)}\,dy_p&=[1-4N_p(1-(\mathcal{C}+m_{pp})(1-\mu_{Aa}-\mu_{aA})-\mu_{aA})]\log(1-y)\\&+[1-4N_p(\mu_{aA}+\mathcal{C}(1-\mu_{Aa}-\mu_{aA}))]\log(y)\\&-[2N_p m_{pp}(1-\mu_{Aa}-\mu_{aA})]\log\left(w_{AA}y^2+2w_{Aa}y(1-y)+w_{aa}(1-y)^2\right)+C,\end{aligned}$$

where $C$ is the constant of integration which cancels out when evaluating the definite integral, and $\mathcal{C}$ is defined as:

$$\mathcal{C}=\sum_{q\neq p}m_{pq}\frac{w_{AA}y_q^2+w_{Aa}y_q(1-y_q)}{w_{AA}y_q^2+2w_{Aa}y_q(1-y_q)+w_{aa}(1-y_q)^2}.$$

Therefore, we can rewrite the Chang and Cooper scheme in two distinct steps:

1. For each $j=0,\ldots,N+1$ evaluate $\omega_j$ using Equation S22 and $\delta_j$ using:

$$\delta_j=\begin{cases}\dfrac{1}{\omega_j}+\dfrac{1}{1-\exp(\omega_j)}, & \text{if } \omega_j\neq 0,\\ 0.5, & \text{if } \omega_j=0.\end{cases}$$

2. For each time step $\tau_n$, $n=1,\ldots,M$ obtain $\alpha^{n+1}$ from $\alpha^n$.

$$\alpha^n=\begin{pmatrix}\alpha(\tau_n,y_1)\\ \alpha(\tau_n,y_2)\\ \vdots\\ \alpha(\tau_n,y_N)\end{pmatrix}$$

By substituting the numerical flux (Equation S21) back into the implicit difference operator (Equation S20) we obtain the tridiagonal system. Having the value for $\alpha^n$ on time step $\tau_n$, we obtain the value $\alpha^{n+1}$ on the next level of time grid by solving:

$$(\boldsymbol{I} + \boldsymbol{B})\,\alpha^{n+1} = \alpha^n.$$

where $\boldsymbol{I}$ is the identity matrix and $\boldsymbol{B}$ is defined as follows:

$$\boldsymbol{B} = \begin{pmatrix} -X_1 & -Z_1 & 0 & \dots & 0 \\ X_1 & -X_2 - Z_1 & -Z_2 & \dots & 0 \\ 0 & X_2 & -X_3 - Z_2 & \dots & 0 \\ \vdots & \ddots & \ddots & \ddots & \vdots \\ 0 & \dots & X_{D-2} & -X_{D-1} - Z_{D-2} & -Z_{D-1} \\ 0 & \dots & 0 & X_{D-1} & -Z_{D-1} \end{pmatrix}. \tag{S23}$$

$$X_j = \frac{\Delta\tau_{n+1}}{\Delta_j}\left[\frac{1}{\Delta_{j+1}} C(y_{j+1/2}) - \delta_j B(y_{j+1/2})\right],$$
$$Z_j = \frac{\Delta\tau_{n+1}}{\Delta_j}\left[\frac{1}{\Delta_{j+1}} C(y_{j+1/2}) + (1-\delta_j) B(y_{j+1/2})\right].$$

We iterate this system forwards through the temporal observation sequence, solving the tridiagonal matrix inversion efficiently in $\mathcal{O}(D)$ time using the Thomas algorithm used in Gutenkunst et al. (2009).

#### 2.3 1D Backward Equation (Crank-Nicolson Scheme)

As an alternative way to calculate the likelihood for temporal HMM, we can run backward pass, which requires solving backward Kolmogorov equation. We also start with the case of one population and neglect population and time segment indices for notation simplicity. Therefore, we denote the target conditional probability function as $\beta(\tau, x)$ and we aim to solve:

$$\frac{\partial}{\partial\tau}\beta(\tau,x) = \frac{1}{2}b(x)\frac{\partial^2}{\partial x^2}\beta(\tau,x) + a(x)\frac{\partial}{\partial x}\beta(\tau,x),$$
$$\text{with a boundary condition: } \frac{\partial}{\partial x}\beta(\tau,x) = 0, \quad \text{at } x = 0,\ x = 1.$$

Because the boundaries at $x = 0$ and $x = 1$ act as reflecting barriers, they result in $\beta(\tau, x_0) = \beta(\tau, x_1)$ and $\beta(\tau, x_{D+1}) = \beta(\tau, x_D)$.

To numerically evaluate this system, we discretize using the standard Crank-Nicolson finite difference scheme. We denote $\beta_j^n = \beta(\tau_n, x_j)$ and use the following approximations for the first and second derivatives:

$$\frac{\partial}{\partial t}\beta_j^n \approx \frac{\beta_j^{n+1} - \beta_j^n}{\Delta t},$$
$$\frac{\partial}{\partial x}\beta_j^n \approx \frac{\beta_{j+1}^n - \beta_{j-1}^n}{\Delta_j + \Delta_{j+1}},$$
$$\frac{\partial^2}{\partial x^2}\beta_j^n \approx \frac{\beta_{j+1}^n - \beta_j}{\frac{1}{2}\Delta_{j+1}(\Delta_j + \Delta_{j+1})} - \frac{\beta_j^n - \beta_{j-1}^n}{\frac{1}{2}\Delta_j(\Delta_j + \Delta_{j+1})}.$$

Let $\beta^n$ denote the column vector representing the conditional probabilities evaluated at the $D$ interior grid points at time $\tau_n$:

$$\beta^n = \begin{pmatrix} \beta(\tau_n, x_1) \\ \beta(\tau_n, x_2) \\ \vdots \\ \beta(\tau_n, x_N) \end{pmatrix}$$

By applying proposed approximations and boundary conditions to our backward equation, the solution dynamic is captured by a tridiagonal linear system:

$$(\boldsymbol{I} - \boldsymbol{A})\, \beta^{n+1} = (\boldsymbol{I} + \boldsymbol{A})\, \beta^{n}.$$

with matrix $\boldsymbol{A}$ defined as:

$$\boldsymbol{A} = \begin{pmatrix} -A_1 & A_1 & 0 & \dots & 0 \\ C_2 & -A_2 - C_2 & A_2 & \dots & 0 \\ 0 & C_3 & -A_3 - C_3 & \dots & 0 \\ \vdots & \ddots & \ddots & \ddots & \vdots \\ 0 & \dots & C_{D-1} & -A_{D-1} - C_{D-1} & A_{D-1} \\ 0 & \dots & 0 & C_D & -C_D \end{pmatrix}. \tag{S24}$$

$$A_j = \frac{1}{4}\left(\Delta_j \lambda_j^b + \lambda_j^a\right),$$
$$C_j = \frac{1}{4}\left(\Delta_{j+1} \lambda_j^b - \lambda_j^a\right).$$
$$\lambda_j^b = b(x_j)\frac{2\Delta\tau_{n+1}}{\Delta_j \Delta_{j+1}(\Delta_j + \Delta_{j+1})},$$
$$\lambda_j^a = a(x_j)\frac{2\Delta\tau_{n+1}}{\Delta_j + \Delta_{j+1}}.$$

We iterate this system backwards through the temporal observation sequence, solving the tridiagonal matrix inversion efficiently in $\mathcal{O}(D)$ time using the Thomas algorithm.

#### 2.4 Extension to Multidimensional Case (Alternating Direction Method)

Having derived the one-dimensional numerical schemes for both the forward and backward passes, we now extend our approach to $P$-dimensional partial differential equations (representing $P$ interacting populations). For a spatial grid with $D$ interior nodes per dimension evaluated over $M$ total time steps, solving a fully coupled implicit multidimensional scheme directly requires flattening the state space into a massive banded matrix. This results in an intractable time complexity of $\mathcal{O}(MD^{3P-2})$ and a memory complexity of $\mathcal{O}(D^{2P-1})$.

To avoid this computational cost, we utilize an alternating direction method based on operator splitting (Press, 2007, p. 1051). By decomposing the multidimensional scheme into a sequence of easily solvable 1D tridiagonal systems, this approach drastically reduces the time complexity to $\mathcal{O}(MD^P)$ and the memory requirements to just $\mathcal{O}(D^P)$.

Let $u^n$ represent the probability state vector at time $\tau_n$ (which can be either the forward probability density $\alpha^n$ or the backward conditional probability $\beta^n$). Any of our one-dimensional numerical schemes for dimension $p$ can be expressed in the following general matrix form:

$$\boldsymbol{C}_p u^{n+1} = \boldsymbol{D}_p u^n.$$

The exact definitions of the matrices $\boldsymbol{C}_p$ and $\boldsymbol{D}_p$ depend on the chosen numerical scheme. They are built using the respective tridiagonal operator matrices $\boldsymbol{B}_p$ (derived for the forward pass in Eq. S23) and $\boldsymbol{A}_p$ (derived for the backward pass in Eq. S24). Crucially, the subscript $p$ indicates that these matrices are defined by the specific drift function $a_p(\mathbf{y})$ and diffusion function $b_{pp}(\mathbf{y})$ governing population $p$. For a full time step $\Delta\tau$, the schemes are defined as:

- **Forward pass (Chang-Cooper):** $\boldsymbol{C}_p = \boldsymbol{I} + \boldsymbol{B}_p$ and $\boldsymbol{D}_p = \boldsymbol{I}$.
- **Backward pass (Crank-Nicolson):** $\boldsymbol{C}_p = \boldsymbol{I} - \boldsymbol{A}_p$ and $\boldsymbol{D}_p = \boldsymbol{I} + \boldsymbol{A}_p$.

Let $\mathbf{i}_{-p} = (i_1, \dots, i_{p-1}, \cdot, i_{p+1}, \dots, i_P)$ denote the spatial grid coordinates omitting the $p$-th dimension. The notation $[\cdot]_{\mathbf{i}_{-p}}$ represents a single one-dimensional "column" extracted along the axis of population $p$.

In our multidimensional model, the drift and diffusion terms are coupled; $a_p$ and $b_{pp}$ depend not only on frequency in population $p$ but also on the frequencies of other populations due to migration. However, along any specific column $\mathbf{i}_{-p}$, the frequencies of all other populations are fixed constants. This allows the multidimensional functions $a_p$ and $b_{pp}$ to be evaluated strictly as 1D functions of $y_p$. Consequently, the 1D operator matrices are uniquely constructed for each individual column. To reflect this explicit spatial dependency, we denote these column-specific matrices as $\boldsymbol{C}_{p,\mathbf{i}_{-p}}$ and $\boldsymbol{D}_{p,\mathbf{i}_{-p}}$.

By splitting the operator over $P$ dimensions, we advance the solution from time level $n$ to $n+1$ via $P$ sequential fractional steps. Applied to our matrix formulation, each spatial operator is implicitly evaluated over the full time step $\Delta\tau$, yielding the following sequence of $P$ equations:

$$\begin{aligned}
\boldsymbol{C}_{1,\mathbf{i}_{-1}} \left[u^{n+1/P}\right]_{\mathbf{i}_{-1}} &= \boldsymbol{D}_{1,\mathbf{i}_{-1}} \left[u^{n}\right]_{\mathbf{i}_{-1}}, \quad \text{for each column } \mathbf{i}_{-1}, \\
\boldsymbol{C}_{2,\mathbf{i}_{-2}} \left[u^{n+2/P}\right]_{\mathbf{i}_{-2}} &= \boldsymbol{D}_{2,\mathbf{i}_{-2}} \left[u^{n+1/P}\right]_{\mathbf{i}_{-2}}, \quad \text{for each column } \mathbf{i}_{-2}, \\
&\vdots \\
\boldsymbol{C}_{P,\mathbf{i}_{-P}} \left[u^{n+1}\right]_{\mathbf{i}_{-P}} &= \boldsymbol{D}_{P,\mathbf{i}_{-P}} \left[u^{n+(P-1)/P}\right]_{\mathbf{i}_{-P}}, \quad \text{for each column } \mathbf{i}_{-P},
\end{aligned}$$

where the fractional superscripts (e.g., $u^{n+1/P}$) represent intermediate algorithmic states rather than physical time steps.

To prevent directional bias and maintain formal temporal accuracy, the sequence in which the 1D directional operators are applied is alternated between consecutive time steps. During the first time step, the spatial dimensions are evaluated in standard ascending order $(1, 2, \ldots, P)$. In the subsequent time step, the evaluation order is exactly reversed $(P, P-1, \ldots, 1)$. This effectively symmetrizes the splitting error.

### 3 Default Simulation and `SweepLink` Configurations

Unless otherwise noted, all simulated benchmarks in this study share a common default configuration, described here once and referenced throughout the main text. Data are generated using the forward-in-time simulator SLiM v4 (Haller and Messer, 2023) and are designed to mimic empirical human data. Each simulated dataset consists of six short chromosomes of equal length, 0.1 Mb. Three chromosomes contain only neutral mutations. The remaining three chromosomes each carry a single hard selective sweep introduced at the midpoint of the chromosome, with selection coefficients $s = 0.01$ (weak), $s = 0.02$ (moderate), and $s = 0.05$ (strong), respectively. Population parameters are chosen to resemble realistic human populations. The effective population size is set to $N = 10{,}000$ individuals (Takahata, 1993). The mutation rate is $\mu = 10^{-8}$ per site per generation. The recombination rate is $r = 10^{-8}$ cM per Mb. Sampling begins once the selected allele reaches a frequency of 30%. Genetic data are sampled at $K = 11$ consecutive time points, with $\Delta t = 16$ generations between consecutive samples, spanning a total of 160 generations. This sampling regime mirrors the real human data analyzed in this study, which includes samples dating back approximately 4,000 years before present. At each time point, genetic data are sampled for $n = 25$ diploid individuals. This entire simulation setup is replicated 100 independent times per experimental configuration.

Unless stated otherwise, `SweepLink` is launched with quadratic grid with 50 grid points for the numerical scheme of the diffusion equation. The choice of quadratic grid over other implemented grid types is defined below. The following grid on positive values of selection is used (13 states): 0.0, 0.005, 0.010, 0.015, 0.020, 0.025, 0.030, 0.035, 0.040, 0.045, 0.050, 0.055, 0.060. The dominance coefficient is fixed to 0.5 (no dominance).

### 4 Data filtering and Optimal Grid Type

#### 4.1 Experimental Setup

We explore the mean likelihood surfaces for population size $N$ and selection $s$ to evaluate how temporal layer of `SweepLink` depends on choice of spatial grid point distribution and on data filtering. Using the default simulation configuration described in Section 3, we compare four implemented spatial grid types: uniform, quadratic, Chebyshev, and logistic (Figure S3).

To assess the estimation of $N$, we constructed expected likelihood surfaces across sample sizes of $n = 10, 25, 100$, and 500 using 100 simulated datasets, each comprising 300 linked neutral loci. Next, to evaluate the inference of $s$, we generated expected likelihood surfaces for a representative sample size of $n = 25$ using 100 simulated datasets, each containing 500 unlinked selective sweeps with varying selection strengths ($s = 0.0, 0.01, 0.02$, and 0.05).

We tested `SweepLink` using four grid types for the allele frequency: uniform, quadratic, Chebyshev, and logistic (Figure S3). We evaluated the shapes of the resulting likelihood surfaces to determine which grid type provides the most accurate and reliable parameter estimation. Additionally, we compared the performance of `SweepLink` against `ApproxWF` and `diplo-locus`.

#### 4.2 Results

Figure S4 illustrates the obtained expected likelihood surfaces for the population size ($N$) evaluated at sample sizes of $n = 10, 25, 100$, and 500.

The topology of the likelihood surface for population size estimation is strongly influenced by the sample size ($n$). In small-sample regimes ($n = 10, 25$), the likelihood increases sharply for small population sizes but reaches a plateau near the ground truth, failing to decay for larger parameter values. This indicates that while small sample sizes can provide a lower bound for population size, they lack the resolution to define an upper bound. In contrast, larger sample sizes ($n = 100, 500$) produce a clearly unimodal likelihood surface with a distinct peak at the ground truth. We note that `SweepLink` with the uniform grid appears to be the best choice in most cases, especially in small-sample regimes ($n = 10, 25$), as it maintains a small likelihood peak near the true value before the plateau. In the large-sample regime ($n = 500$) all grid types perform well. Across all scenarios, `SweepLink` with the uniform grid provides more accurate maximum likelihood estimates than `ApproxWF`.

The expected likelihood surfaces for the selection coefficient ($s$) for a representative sample size of $n = 25$ are shown on Figure S5.

The topology of the likelihood surface for $s$ is moderately influenced by the choice of grid type. The most well-defined, unimodal surfaces were observed with the uniform grid, followed closely by the Chebychev and quadratic grids. This makes uniform grid a robust default grid type for the joint inference. In contrast, the tanh and logistic grids produced flatter likelihood profiles with reduced curvature. We note that this ordering of grid performance directly corresponds to the density of grid nodes at intermediate allele frequencies (around 0.5). While the uniform grid maintains a constant resolution across the entire frequency spectrum, the quadratic, Chebyshev, tanh, and logistic grids progressively deplete the number of nodes in the central region in favor of increasingly dense clustering near the boundaries (Figure S3). Because the deterministic effect of selection drives the largest allele frequency shifts at intermediate frequencies, excessively sparse grids in this region fail to adequately resolve the selection dynamics, resulting in the observed loss of likelihood curvature.

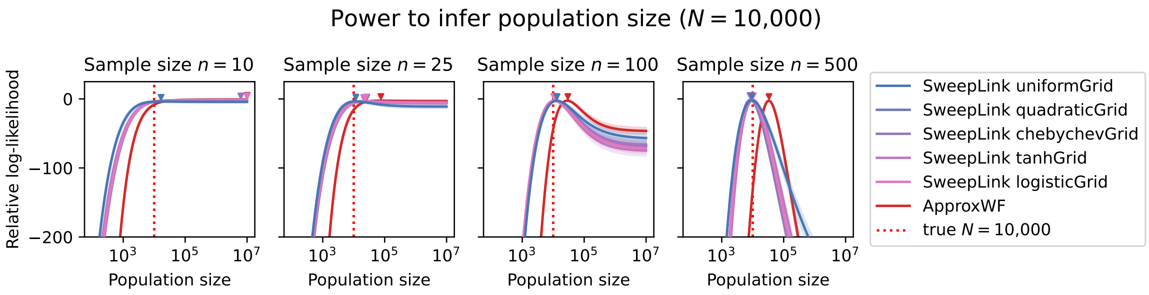

Figure S4: Mean relative likelihood surfaces for population size estimation using `SweepLink` across various grid types for allele frequencies and `ApproxWF`. Solid lines represent the mean and shaded regions denote the confidence intervals calculated across 100 independent datasets, each containing 300 loci. The triangle marks the maximum point of the mean likelihood. The ground truth population size of 10,000 individuals is indicated by the vertical red dashed line.

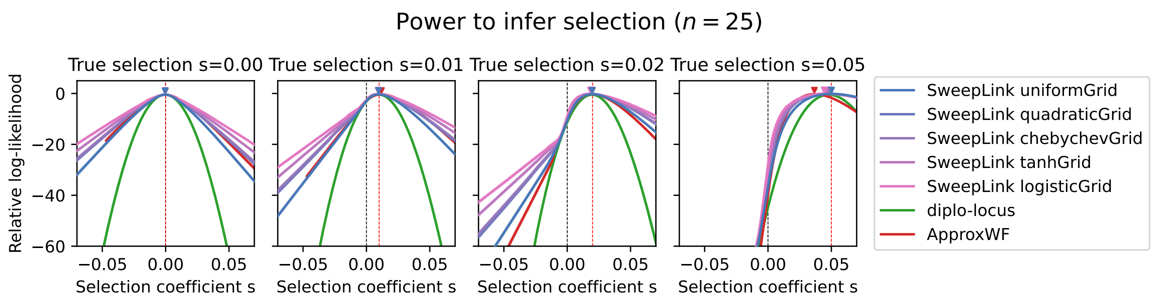

Figure S5: Mean relative likelihood surfaces for selection estimation using `SweepLink` across various grid types, `ApproxWF` and `diplo-locus`. Solid lines represent the mean and shaded regions denote the confidence intervals calculated across 100 independent datasets, each containing 500 unlinked loci under selection. The triangle marks the maximum point of the mean likelihood. The ground truth selection coefficients are indicated by the vertical red dashed line.

The precision of the selection estimates is sensitive to the underlying intensity of selection. For neutrality ($s = 0$) and moderate selection coefficients ($s = 0.01$ and $s = 0.02$), `SweepLink` produces well-behaved likelihood surfaces with clear peaks centered at the ground truth. Under strong selection ($s = 0.05$), the surface exhibits a plateau at higher parameter values, suggesting a lack of statistical resolution to define an upper bound. Crucially, however, even when the surface is nearly flat, `SweepLink` consistently maintains a maximum likelihood estimate very close to the true value. This indicates that while the confidence in the exact value may decrease under strong selection, the point estimate remains highly accurate. In comparison with other methods, `diplo-locus` produces highly resolved likelihood surfaces with sharp peaks near the ground truth. However, it must be noted that `diplo-locus` requires the population size to be known a priori, whereas `SweepLink` performs a joint inference of both $N$ and $s$. While `ApproxWF` demonstrated likelihood shapes similar to—and occasionally steeper than—`SweepLink`, it exhibited a significant downward bias under the strong selection ($s = 0.05$) scenario.

### 5 Benchmarking Against Competitor Single-Locus Tools

Table S1: **Locus-level detection performance.** False-positive rate and true positive rate by true selection coefficient $s$, evaluated per locus (no window-based grouping) at each tool's own best-MCC threshold. `SweepLink` attains the highest overall detection power and the highest power at weak and moderate selection ($s = 0.01$, $s = 0.02$) among all tools, but also by far the highest false-positive rate, reflecting spatially correlated false calls produced by its linkage-aware model rather than a genuine excess of erroneous detections (see main text for the window-based comparison).

| | | | True Positive Rate by selection coefficient $s$ | | | | | |
|---|---|---|---|---|---|---|---|---|
| **Tool** | **FPR** | **Overall TPR** | **0.01** | **0.02** | **0.03** | **0.04** | **0.05** | **MCC** |
| `SweepLink` | 0.052% | **81%** | 14% | 90% | 100% | 100% | 100% | 0.838 |
| `ApproxWF` | 0.010% | 73% | 3% | 63% | 99% | 100% | 100% | 0.843 |
| `diplo-locus` | **0.004%** | 75% | 3% | 73% | 100% | 100% | 100% | **0.861** |
| `bmws` | 0.017% | 55% | 0% | 7% | 70% | 97% | 100% | 0.714 |

### 6 Extended Benchmarking of `SweepLink` on Simulated Data

#### 6.1 Experimental Setup

We further validate the robustness of the `SweepLink` framework under diverse evolutionary and sampling scenarios. We first establish the default configuration, which serves as a baseline for all subsequent experiments. To assess the impact of individual characteristics on the accuracy of `SweepLink`, we varied one characteristic at a time—for example, sample size, sequence length or recombination rate—while keeping all other parameters constant at their default values. This approach enables us to quantify how each parameter influences the accuracy of inference.

For each experiment, we generated six chromosomes, each with a length of 0.1 Mb in the default configuration. Three chromosomes contained only neutral mutations. The remaining three chromosomes had a hard selective sweep introduced at the middle position, with a selection coefficient equal to 0.01, 0.02, or 0.05, respectively. Population parameters were chosen to resemble realistic human populations: population size was set to 10,000 individuals, mutation rate to $10^{-8}$ per site per generation, and recombination rate to $10^{-8}$ cM per Mb. Sampling was performed once the selected allele reached a frequency of 30%. Genetic data were sampled at eleven consecutive time points, with 16 generations between each, thereby spanning 160 generations. This sampling regime reflects the real human data analyzed in this study, which includes samples dating approximately 4,000 years before present. At each time point, genetic data for 25 diploid individuals were sampled. Sweeplink is launched with 50 grid points for the numerical scheme of the diffusion equation. The following grid on positive values of selection is used (13 states): 0.0, 0.005, 0.010, 0.015, 0.020, 0.025, 0.030, 0.035, 0.040, 0.045, 0.050, 0.055, 0.060. The dominance coefficient is fixed to 0.5 (no dominance).

In summary, we test `SweepLink` across the following characteristics: 1) sample size, 2) sequence length, 3) number of time points, 4) binning size, 5) starting allele frequency of the selective sweep, 6) recombination rate, and 7) grid size in the numerical scheme for the diffusion equation.

For the sample size experiments, the number of sampled diploid individuals was kept equal at all time points, with values of 5, 10, 25, 50, 100, 250, or 500. For sequence length variation, all six chromosomes were simulated with the same length, using one of the following values in separate experiments: 0.05, 0.1, 0.2, 0.5, or 1 Mb. For experiments varying the number of time points, we used 6, 11, 21, and 41 time points, spanning uniformly over 160 generations.

In order to assess the effect of binning, we sampled genetic data from two randomly chosen diploid individuals from the population at every generation over a total span of 160 generations. We defined bins by dividing this time span into intervals of equal length, and used the midpoint of each interval as the representative time point for that bin. The interval lengths tested were 2, 4, 8, 16, 32, 40, and 80 generations, which correspond to 80, 40, 20, 10, 5, 4, and 2 bins, respectively. For each bin, all samples from generations falling within the interval were assigned to the bin represented by that interval's midpoint. At the boundaries between adjacent intervals, one sample from that generation was assigned to one bin and the other sample to the next bin. For example, with an interval length of 40 generations, the bins cover the generation ranges 0–20, 20–60, 60–100, 100–140, and 140–160 (relative to the first sampling point), with representative time points at 0, 40, 80, 120, and 160 generations after the first sampling point. The central bins each contain an equal number of samples, whereas the two edge bins (at the first and last time points) may include fewer samples, as they are only one-sided. This procedure ensures that the binning scheme is as balanced as possible, except at the boundaries of the interval.

In a separate set of experiments, we investigate how varying the initial frequency of the derived allele at the selected locus influences inference results. To do this, we initiate sampling at the generation when the frequency of the derived allele undergoing a hard sweep first reached one of the following values: 0.01, 0.1, 0.2, 0.3, 0.4, 0.5, 0.6, 0.7, 0.8, or 0.9.

The linkage between loci and the magnitude of the hitch-hiking effect are influenced by the recombination rate. While SweepLink estimates the recombination rate automatically, we expect its inference accuracy to be higher under stronger linkage, that is, at lower recombination rates. To explore this, we varied the recombination rate across five values: $10^{-9}$, $10^{-8}$, $10^{-7}$, $10^{-6}$ and $10^{-5}$ cM per Mb.

The final set of experiments assessed how the grid size in the numerical scheme for the diffusion equation in SweepLink affects inference accuracy. We tested grid sizes of 5, 10, 20, 50, and 100 points.

#### 6.2 Results

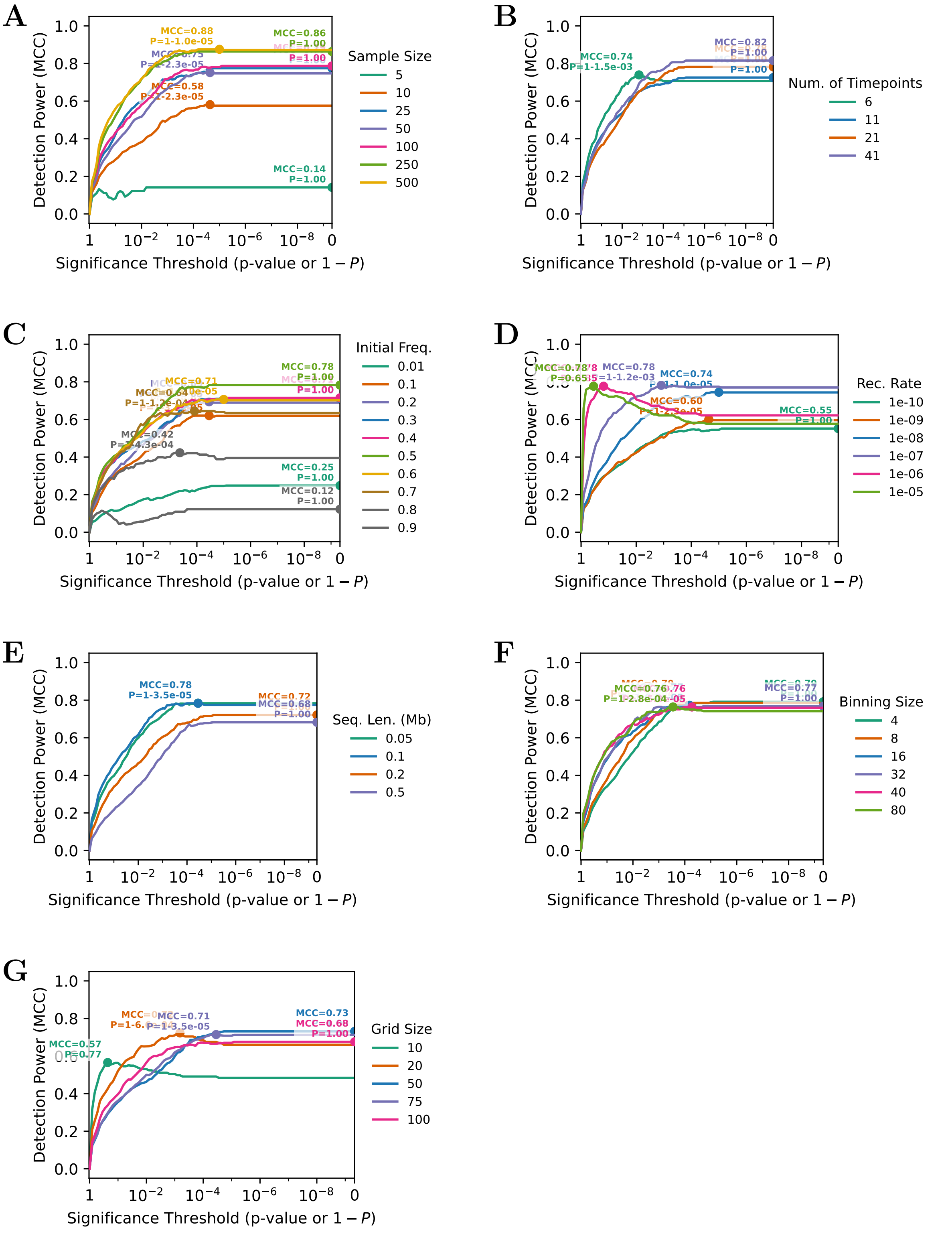

Figure S6: **Detection power across varied parameters and study designs.** (A) Impact of varying the number of diploid samples per timepoint. (B) Impact of the number of temporal sampling points. (C) Effect of the initial allele frequency of selected variant at the onset of sampling. (D) Robustness of the model to the number of linked loci included in the regional genomic context. (E) Sensitivity to varying background recombination rates. (F) Impact of spatial/temporal data binning strategies. (G) Effect of the internal computational grid resolution.

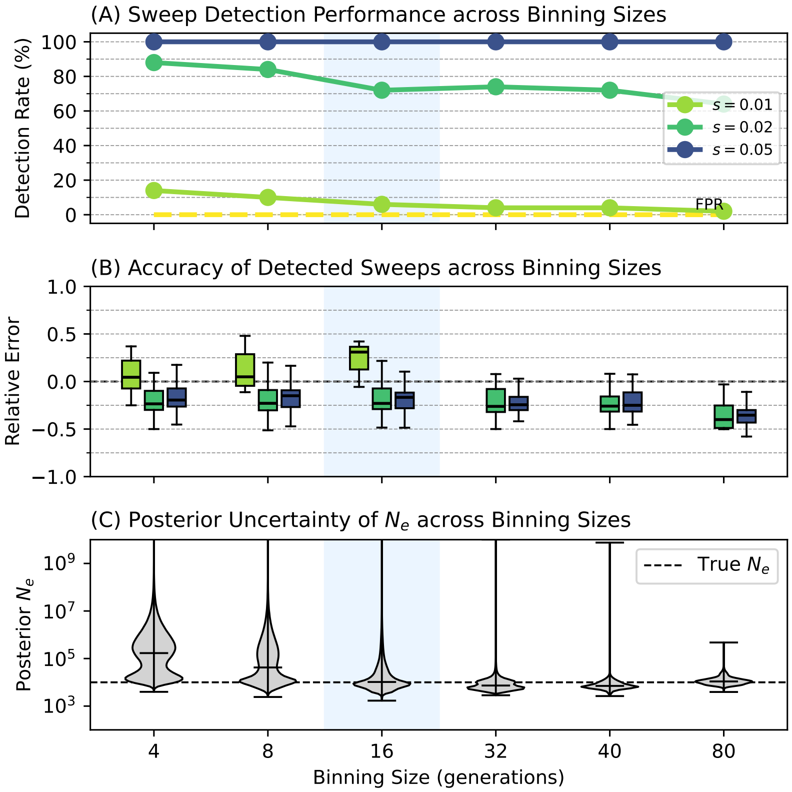

Figure S7

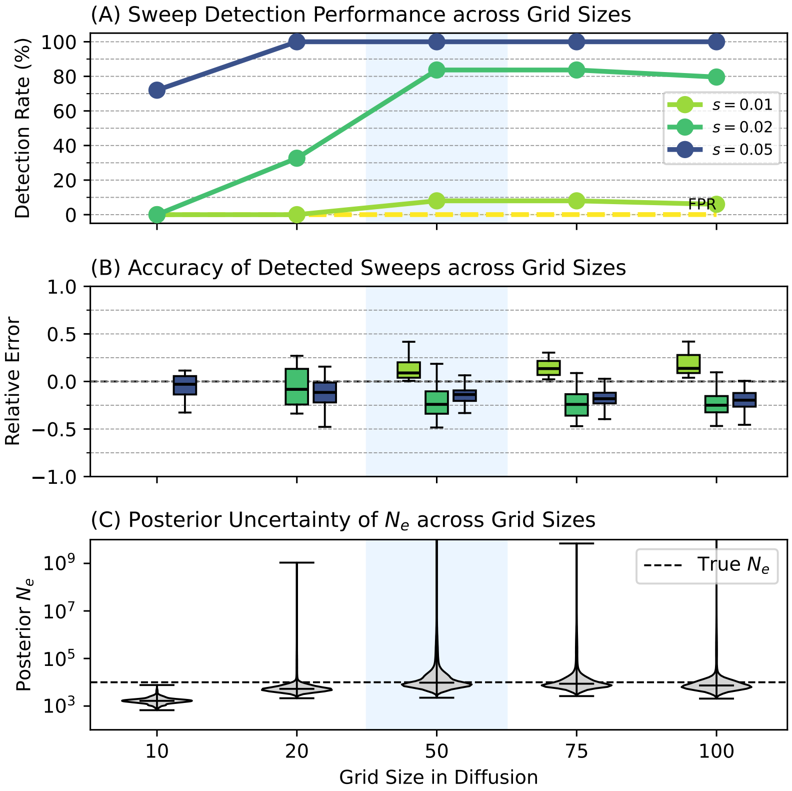

Figure S8

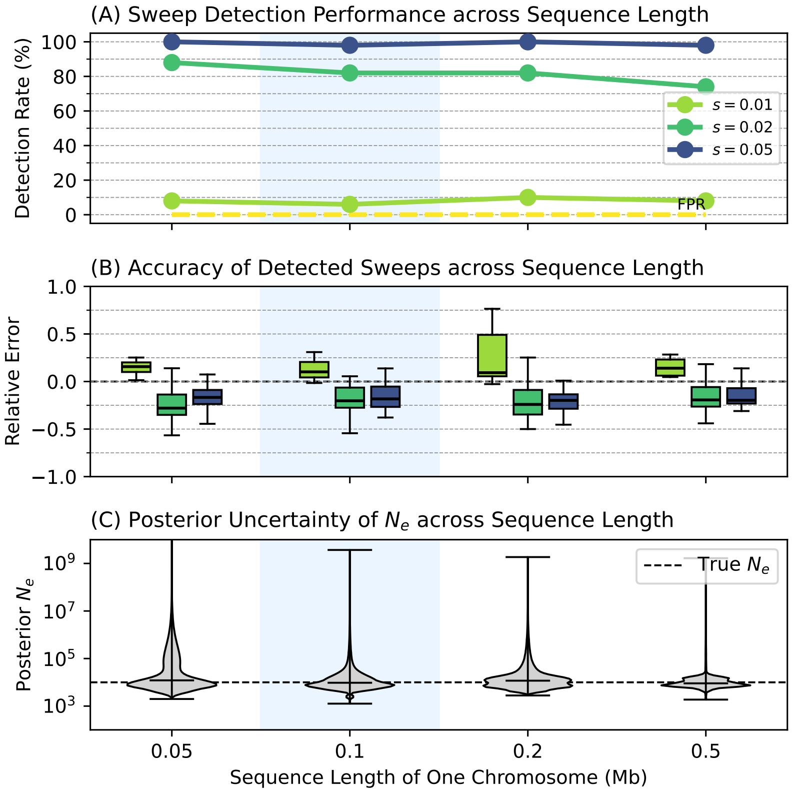

Figure S9

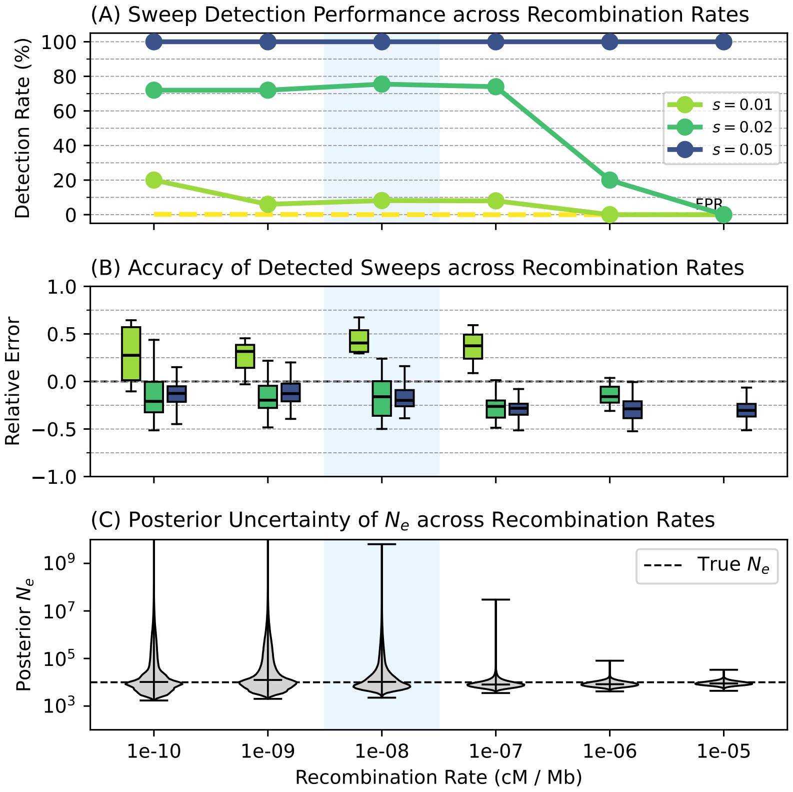

Figure S10

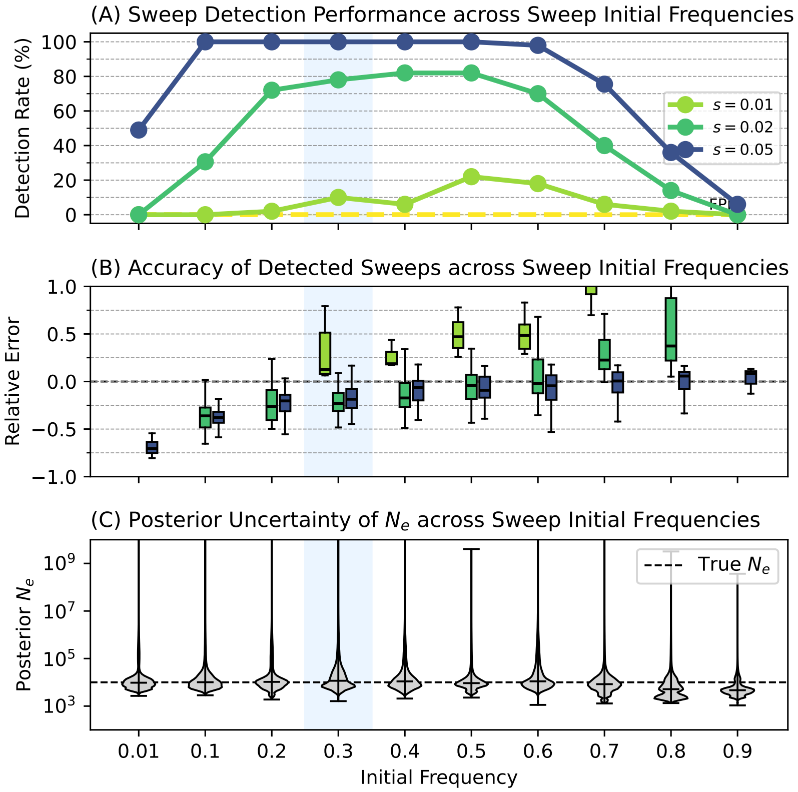

Figure S11

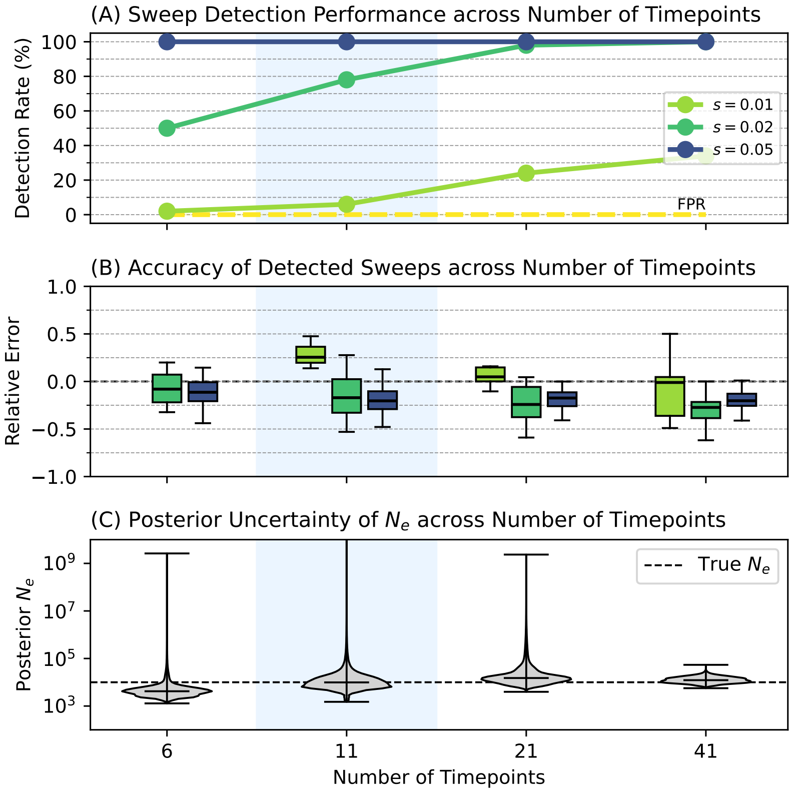

Figure S12

### 7 Results for British data

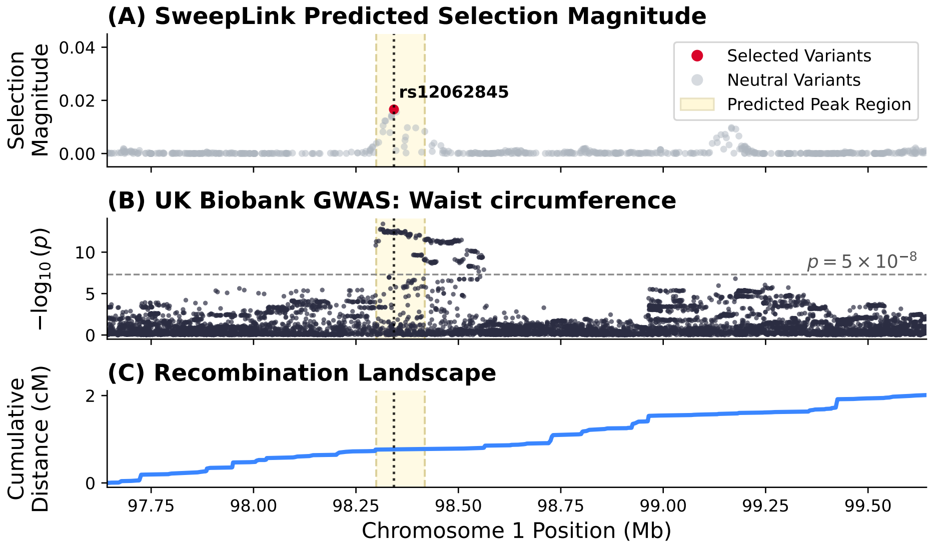

Figure S13: **Genomic landscape of the selection signal at the DPYD locus on chromosome 1.** (A) Predicted selection magnitudes generated by `SweepLink`. Red points indicate variants classified as target variants under selection ($P = 1.0$), while gray points represent neutral variants. The focal variant, rs12062845, is marked by a vertical dotted line. (B) Local Manhattan plot of GWAS summary statistics from the UK Biobank (Neale lab) for the trait "Waist circumference" (OpenGWAS ID: ukb-b-9405). This trait colocalizes with predicted selection signal (colocalization probability = 0.98). The horizontal dashed line denotes the standard genome-wide significance threshold ($p = 5 \times 10^{-8}$). (C) Cumulative recombination landscape (cM) based on the 1000 Genomes Project genetic map for the GBR population.

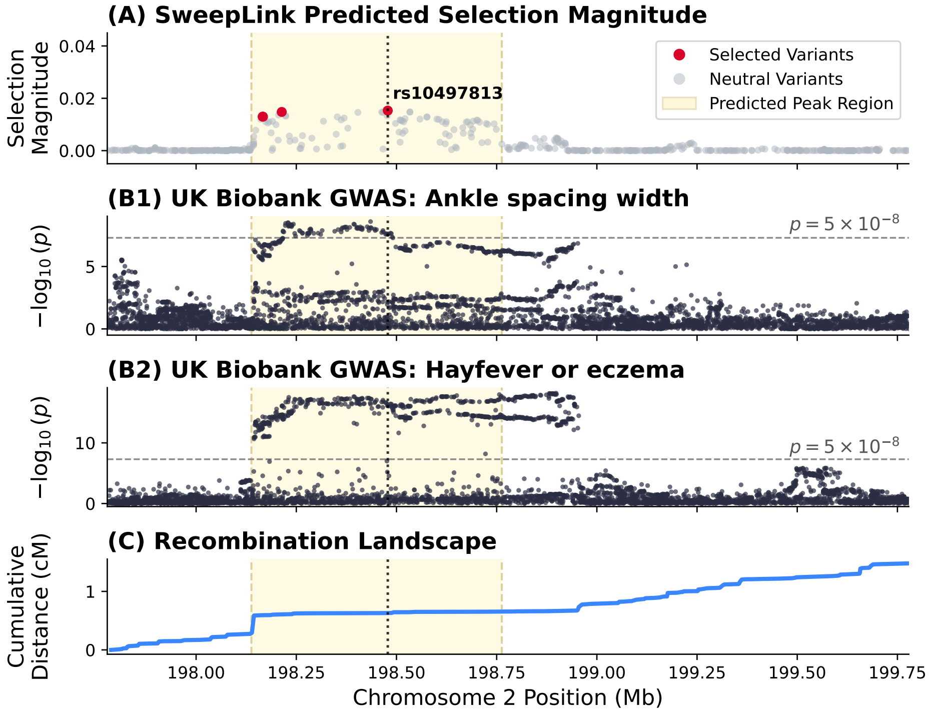

Figure S14: **Genomic landscape of the selection signal at the RFTN2 locus on chromosome 2.** (A) Predicted selection magnitudes generated by `SweepLink`. Red points indicate variants classified as target variants under selection ($P = 1.0$), while gray points represent neutral variants. The focal variant, rs1455653, is marked by a vertical dotted line. (B1) Local Manhattan plot of GWAS summary statistics from the UK Biobank (Neale lab) for the trait "Ankle spacing width" (OpenGWAS ID: ukb-b-4080). This trait colocalizes with predicted selection signal (colocalization probability = 0.97). The horizontal dashed line denotes the standard genome-wide significance threshold ($p = 5 \times 10^{-8}$). (B2) Local Manhattan plot of GWAS summary statistics from the UK Biobank (Neale lab) for the trait "... diagnosed by doctor: Hayfever, allergic rhinitis or eczema" (OpenGWAS ID: ukb-b-17241). This trait colocalizes with predicted selection signal (colocalization probability = 0.84). (C) Cumulative recombination landscape (cM) based on the 1000 Genomes Project genetic map for the GBR population.

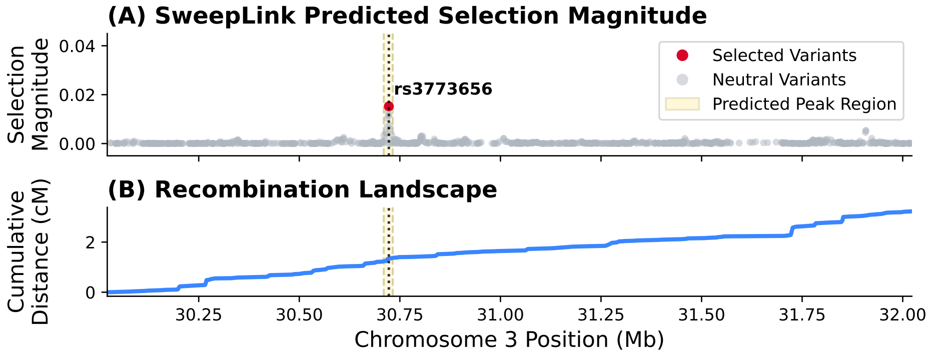

Figure S15: **Genomic landscape of the selection signal at the TGFBR2 locus on chromosome 3.** The focal variant targeted by selection at this locus (rs3773656) maps to a known enhancer region. (A) Predicted selection magnitudes generated by `SweepLink`. Red points indicate variants classified as target variants under selection ($P = 1.0$), while gray points represent neutral variants. (C) Cumulative recombination landscape (cM) based on the 1000 Genomes Project genetic map for the GBR population.

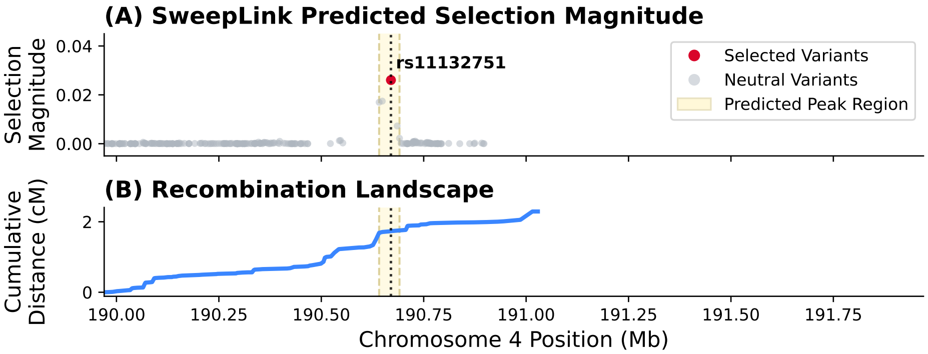

Figure S16: **Genomic landscape of the selection signal at an intergenic region on chromosome 4.** (A) Predicted selection magnitudes generated by `SweepLink`. Red points indicate variants classified as target variants under selection ($P = 1.0$), while gray points represent neutral variants. The focal variant, rs11132751, is marked by a vertical dotted line. (C) Cumulative recombination landscape (cM) based on the 1000 Genomes Project genetic map for the GBR population.

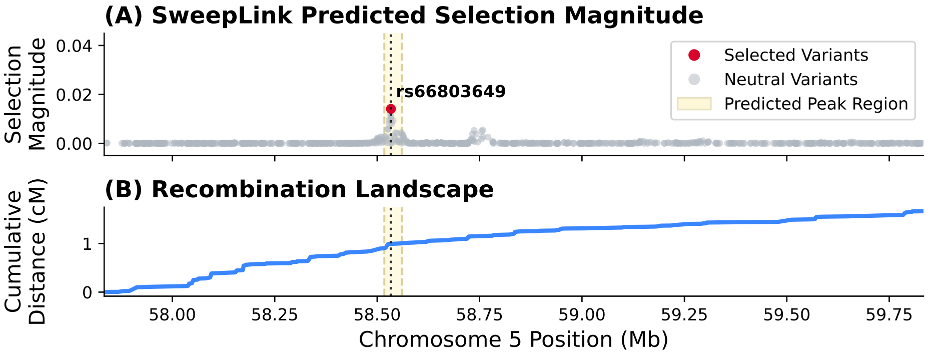

Figure S17: **Genomic landscape of the selection signal at the PDE4D locus on chromosome 5.** (A) Predicted selection magnitudes generated by `SweepLink`. Red points indicate variants classified as target variants under selection ($P = 1.0$), while gray points represent neutral variants. The focal variant, rs66803649, is marked by a vertical dotted line. (C) Cumulative recombination landscape (cM) based on the 1000 Genomes Project genetic map for the GBR population.

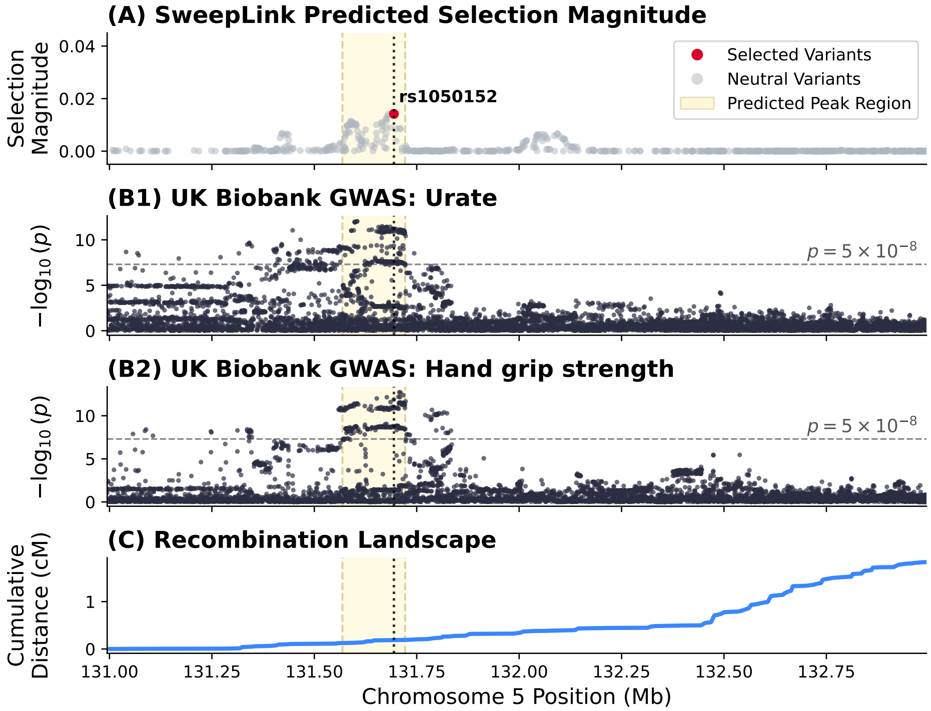

Figure S18: **Genomic landscape of the selection signal at the SLC22A4/5 locus on chromosome 5.** (A) Predicted selection magnitudes generated by `SweepLink`. Red points indicate variants classified as target variants under selection ($P = 1.0$), while gray points represent neutral variants. The focal variant, rs1050152, is marked by a vertical dotted line. (B1) Local Manhattan plot of GWAS summary statistics from the UK Biobank (Neale lab) for the trait "Urate" (OpenGWAS ID: ukb-d-30880_irnt). This trait colocalizes with predicted selection signal (colocalization probability = 0.98). The horizontal dashed line denotes the standard genome-wide significance threshold ($p = 5 \times 10^{-8}$). (B2) Local Manhattan plot of GWAS summary statistics from the UK Biobank (Neale lab) for the trait "Hand grip strength (left)" (OpenGWAS ID: ukb-b-7478). This trait colocalizes with predicted selection signal (colocalization probability = 0.92). (C) Cumulative recombination landscape (cM) based on the 1000 Genomes Project genetic map for the GBR population.

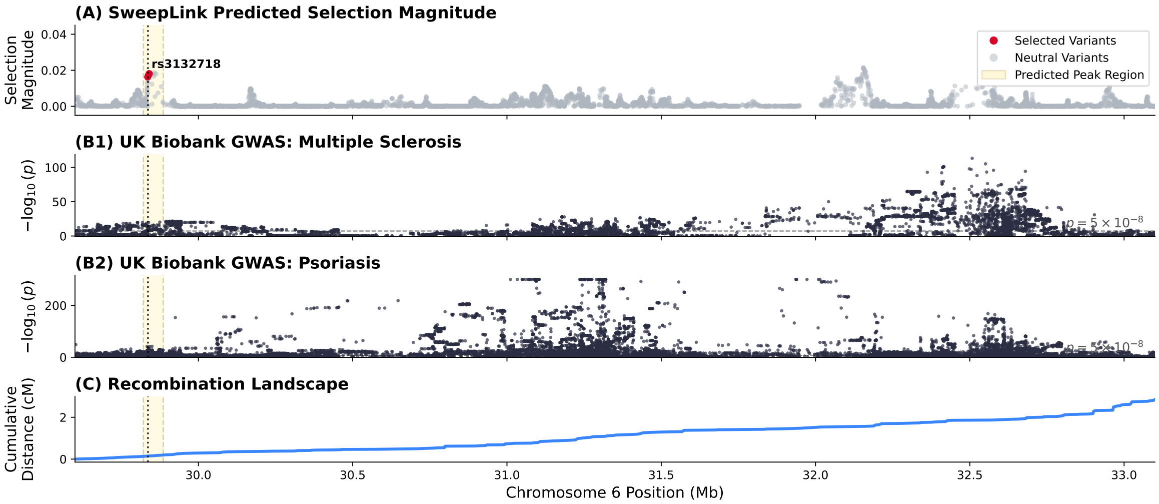

Figure S19: **Genomic landscape of the selection signal at the MHC region on chromosome 6.** No significant colocalization is found for analyzed traits. (A) Predicted selection magnitudes generated by `SweepLink`. Red points indicate variants classified as target variants under selection ($P = 1.0$), while gray points represent neutral variants. The focal variant, rs3132718, is marked by a vertical dotted line. (B1) Local Manhattan plot of GWAS summary statistics from the UK Biobank (Neale lab) for the trait ”Non-cancer illness code, self-reported: multiple sclerosis” (OpenGWAS ID: ukb-b-17670). This trait does not colocalize with predicted selection signal (colocalization probability ¡ 0.01). The horizontal dashed line denotes the standard genome-wide significance threshold ($p = 5 \times 10^{-8}$). (B2) Local Manhattan plot of GWAS summary statistics from the UK Biobank (Neale lab) for the trait ”Non-cancer illness code, self-reported: psoriasis” (OpenGWAS ID: ukb-b-10537). This trait does not colocalize with predicted selection signal (colocalization probability ¡ 0.01). (C) Cumulative recombination landscape (cM) based on the 1000 Genomes Project genetic map for the GBR population.

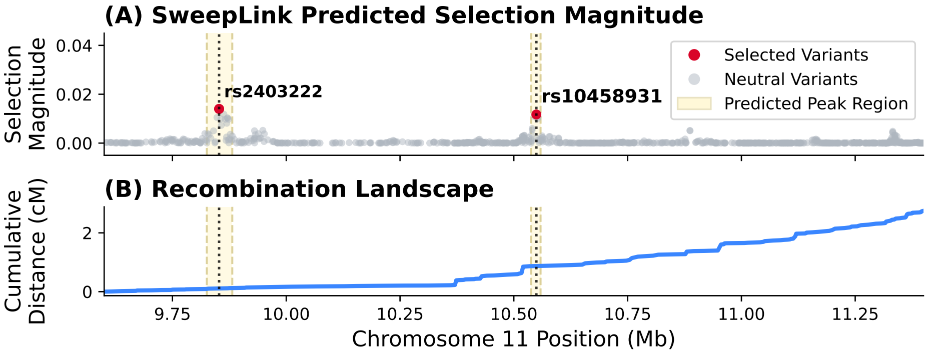

Figure S20: **Genomic landscape of the selection signal at the SBF2 and RNF141 loci on chromosome 11.** (A) Predicted selection magnitudes generated by `SweepLink`. Red points indicate variants classified as target variants under selection ($P = 1.0$), while gray points represent neutral variants. The focal variants, rs2403222 (SBF2) and rs10458931 (RNF141), are marked by a vertical dotted line. (C) Cumulative recombination landscape (cM) based on the 1000 Genomes Project genetic map for the GBR population.

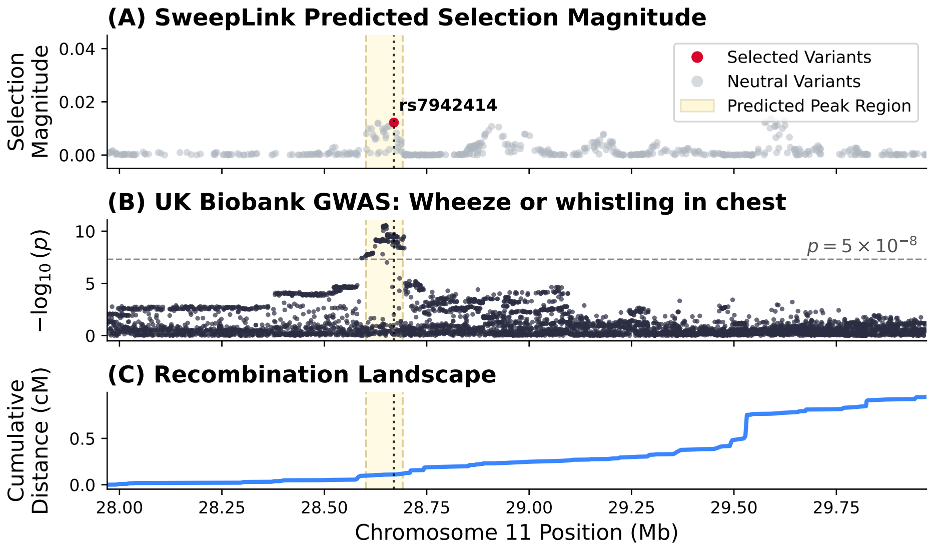

Figure S21: **Genomic landscape of the selection signal at an intergenic region on chromosome 11.** (A) Predicted selection magnitudes generated by `SweepLink`. Red points indicate variants classified as target variants under selection ($P = 1.0$), while gray points represent neutral variants. The focal variant, rs7942414, is marked by a vertical dotted line. (B) Local Manhattan plot of GWAS summary statistics from the UK Biobank (Neale lab) for the trait "Wheeze or whistling in the chest in last year" (OpenGWAS ID: ukb-b-18335). This trait colocalizes with predicted selection signal (colocalization probability = 0.94). The horizontal dashed line denotes the standard genome-wide significance threshold ($p = 5 \times 10^{-8}$). (C) Cumulative recombination landscape (cM) based on the 1000 Genomes Project genetic map for the GBR population.

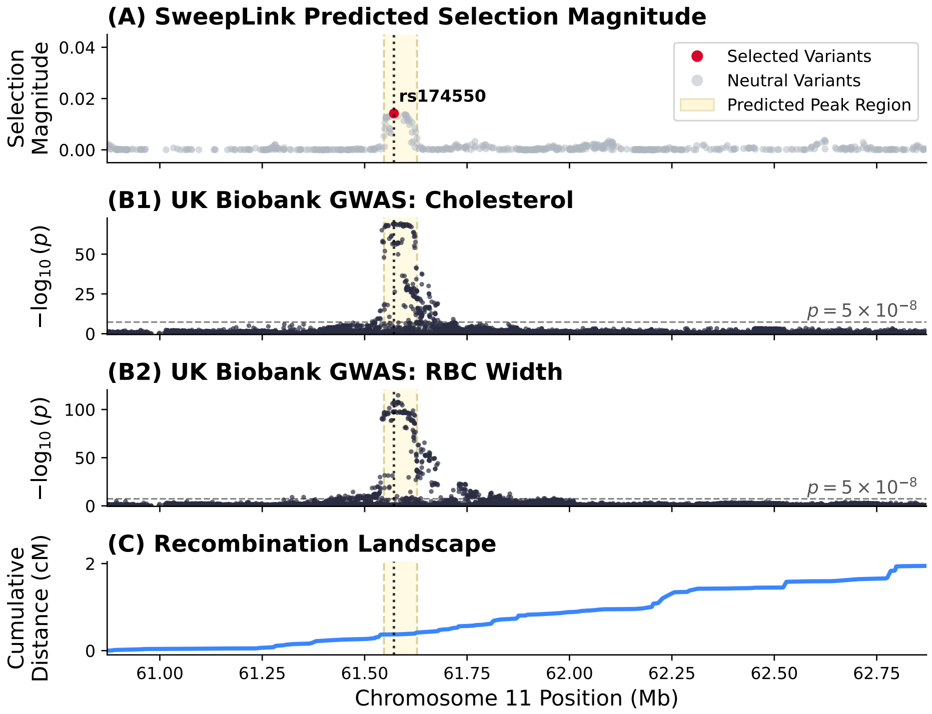

Figure S22: **Genomic landscape of the selection signal at the FADS1/FADS2 region on chromosome 11.** (A) Predicted selection magnitudes generated by `SweepLink`. Red points indicate variants classified as target variants under selection ($P = 1.0$), while gray points represent neutral variants. The focal variant, rs174550, is marked by a vertical dotted line. (B1) Local Manhattan plot of GWAS summary statistics from the UK Biobank (Neale lab) for the trait "Cholesterol" (OpenGWAS ID: ukb-d-30690_raw). This trait colocalizes with predicted selection signal (colocalization probability = 0.99). The horizontal dashed line denotes the standard genome-wide significance threshold ($p = 5 \times 10^{-8}$). (B2) Local Manhattan plot of GWAS summary statistics from the UK Biobank (Neale lab) for the trait "Red blood cell (erythrocyte) distribution width" (OpenGWAS ID: ukb-d-30070_irnt). This trait colocalizes with predicted selection signal (colocalization probability = 0.98). (C) Cumulative recombination landscape (cM) based on the 1000 Genomes Project genetic map for the GBR population.

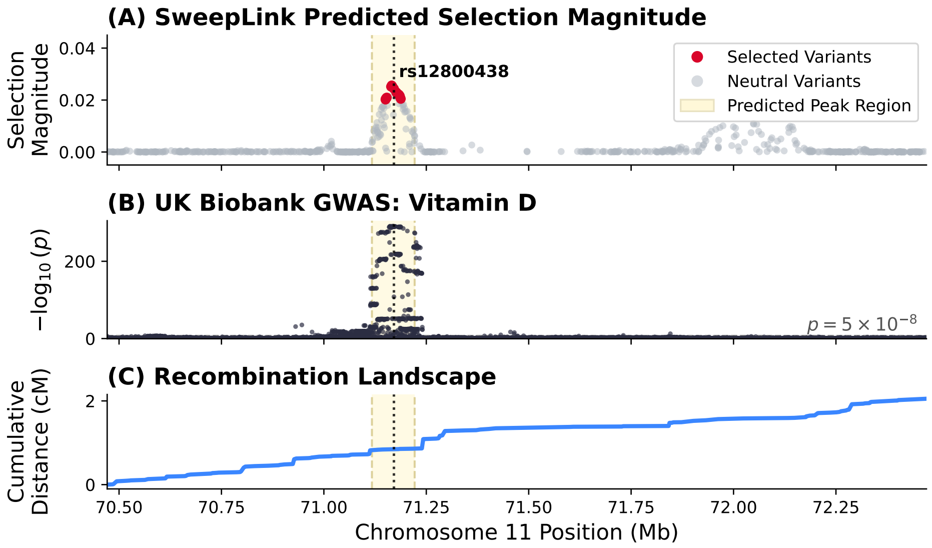

Figure S23: **Genomic landscape of the selection signal at the DHCR7 locus on chromosome 11.** (A) Predicted selection magnitudes generated by `SweepLink`. Red points indicate variants classified as target variants under selection ($P = 1.0$), while gray points represent neutral variants. The focal variant, rs12800438, is marked by a vertical dotted line. (B) Local Manhattan plot of GWAS summary statistics from the UK Biobank (Neale lab) for the trait "Vitamin D" (OpenGWAS ID: ukb-d-30890_irnt). This trait colocalizes with predicted selection signal (colocalization probability = 0.99). The horizontal dashed line denotes the standard genome-wide significance threshold ($p = 5 \times 10^{-8}$). (C) Cumulative recombination landscape (cM) based on the 1000 Genomes Project genetic map for the GBR population.

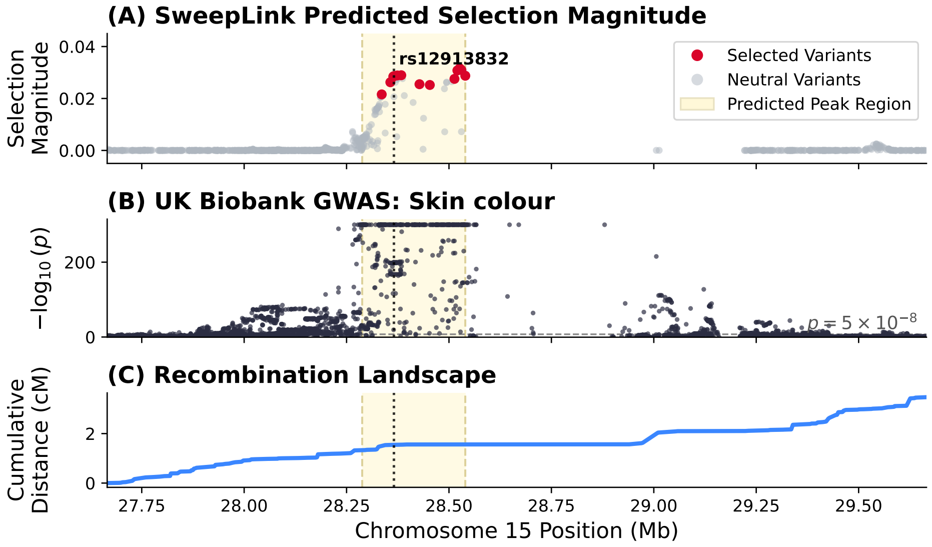

Figure S24: **Genomic landscape of the selection signal at the HERC2 locus on chromosome 15.** (A) Predicted selection magnitudes generated by `SweepLink`. Red points indicate variants classified as target variants under selection ($P = 1.0$), while gray points represent neutral variants. The focal variant, rs12913832, is marked by a vertical dotted line. (B) Local Manhattan plot of GWAS summary statistics from the UK Biobank (Neale lab) for the trait "Skin colour" (OpenGWAS ID: ukb-b-19560). This trait colocalizes with predicted selection signal (colocalization probability = 0.94). The horizontal dashed line denotes the standard genome-wide significance threshold ($p = 5 \times 10^{-8}$). (C) Cumulative recombination landscape (cM) based on the 1000 Genomes Project genetic map for the GBR population.

Samuel Karlin and Howard E Taylor. *A second course in stochastic processes.* Elsevier, 1981.

Anna-Sapfo Malaspinas, Orestis Malaspinas, Steven N Evans, and Montgomery Slatkin. Estimating allele age and selection coefficient from time-serial data. *Genetics*, 192(2):599–607, 2012.

John H Mathews, Kurtis D Fink, et al. *Numerical methods using MATLAB*, volume 4. Pearson prentice hall Upper Saddle River, NJ, 2004.

Keith W Morton and David Francis Mayers. *Numerical solution of partial differential equations: an introduction.* Cambridge university press, 2005.

Lorenzo Pareschi and Mattia Zanella. Structure preserving schemes for nonlinear fokker–planck equations and applications. *Journal of Scientific Computing*, 74(3):1575–1600, 2018.

William H Press. *Numerical recipes 3rd edition: The art of scientific computing.* Cambridge University Press, 2007.

Naoyuki Takahata. Allelic genealogy and human evolution. *Molecular Biology and Evolution*, 10(1): 2–22, 1993.
